# Predictive and error coding for vocal communication signals in the songbird auditory forebrain

**DOI:** 10.1101/2024.02.25.581987

**Authors:** Srihita Rudraraju, Michael E. Turvey, Bradley H. Theilman, Timothy Q. Gentner

## Abstract

Predictive coding posits that sensory signals are compared to internal models, with resulting prediction-error carried in the spiking responses of single neurons. Despite its proposal as a general cortical mechanism, including for speech processing, whether or how predictive coding functions in single-neuron responses to vocal communication signals is unknown. As a proxy internal model, we developed a neural network that uses current sensory context to predict future spectrotemporal features of a vocal communication signal, birdsong. We then represent birdsong as either weighted sets of latent predictive features evolving in time, or as time-varying prediction-errors that reflect the difference between ongoing network-predicted and actual song. Using these spectrotemporal, predictive, and prediction-error song representations, we fit linear/non-linear receptive fields to single neuron responses recorded from caudomedial nidopallium (NCM), caudal mesopallium (CMM) and Field L, analogs of mammalian auditory cortices, in anesthetized European starlings, *Sturnus vulgaris*, listening to conspecific songs. In all three regions, the predictive features of song yield the single best model of song-evoked spiking responses, but unique information about all three representations (signal, prediction, and error) is carried in the spiking responses to song. The relative weighting of this information varies across regions, but in contrast to many computational predictive coding models neither predictive nor error responses are segregated in separate neurons. The continuous interplay between prediction and prediction-error is consistent with the relevance of predictive coding for cortical processing of temporally patterned vocal communication signals, but new models for how prediction and error are integrated in single neurons are required.

## Introduction

Sensory processing of temporally patterned natural signals such as speech and music relies heavily on existing knowledge. One theory for how this knowledge is implemented is “predictive coding” (PC). Put forth originally as a model for visual cortex [1, 2], PC has been proposed as a general cortical mechanism [3, 4] including for speech processing [5, 6]. PC proposes that the brain uses learned input regularities to construct generative models of the world, then tries to ‘fit’ these models by comparison to incoming sensory signals, resulting in a prediction error. When this error goes to zero, the incoming stimulus and the internal model match. In this sense, PC is a form of negative feedback that captures the inferential nature of perception. It is also particularly well-suited to the temporal structure of speech [7, 8, 9, 10, 6] and other complex vocal communication signals such as birdsong, both of which comprise patterned sequential structures with temporal dependencies that stretch across multiple timescales [11]. While several quantitative PC models have been described [1, 3, 12, 13, 14], all share empirical predictions that are largely untested in the context of natural vocal communication signals.

The internal generative models that are thought to guide predictive coding can, in principle, reflect a wide variety of contextual knowledge, including memory and the sensory signal itself [15]. One common framework for studying prediction-error encoding in the auditory system uses stimulus specific adaptation [16], whereby a single stimulus (typically a sine tone) is repeated multiple times leading to a decrease in evoked firing rates. Following this adaption, presentation of a second tone, different in frequency, evokes increased firing rates [17]. While multiple mechanisms can explain repetition suppression [16], the response to the deviant tone is thought to reflect a prediction error [18, 19, 20] potentially via top-down projections [21, 19]. Predictive coding has also been applied in other sensory contexts including learned associations between stimuli either in different sensory modalities or motor-related regions. For example, inputs from the secondary motor cortex suppress neural responses in deep and superficial layers of auditory cortex during self-initiated movements [22, 23], putatively as a mechanism to enhance responses to less predictable sounds. Likewise, learned associations between auditory and visual stimuli can drive experience-dependent suppression within the primary visual cortex [24, 25], and expectation based on olfactory cues can modulate learned responses to auditory stimuli [26]. In humans, predictive coding has been widely applied to explain non-invasive measures of neural responses in a variety of speech and languages studies (e.g., [5]), but we know much less about the underlying neurobiological mechanisms for predictive coding for these kinds of signals.

Songbirds offer a rich domain for investigating the neurobiological mechanisms for predictive coding of vocal communication signals, without having to induce artificial dependencies or associations between arbitrary neutral stimuli. Like speech, the songs of oscine birds are learned during a developmental critical period [27], and they have a spectrally and temporally complex acoustic structure [28] with statistical dependencies that stretch across multiple timescales [11]. Most importantly, the receivers in these communications systems are highly practiced with conspecific vocal signals and the perceptual systems that guide communication behaviors are under heavy selection pressures.

Predictive coding requires a prediction, termed an internal generative model, such as the expectation that a tone will repeat [17] or a movement will create a sound [22]. Here, we hypothesize that a songbird species, European starlings, use the modal statistical structure of their conspecific vocalizations as a default generative model during song perception. To approximate the prediction of this internal model, we train deep neural networks to generate statistical approximations of starling song. Given any song input, this yields a continuous predictive estimate of future vocalizations and a corresponding error relative to true song at each moment in time. We then examine whether and how this generative model functions in the predictive coding of song, by modeling the empirical song-evoked spiking responses of single neurons throughout multiple auditory forebrain regions to determine the information that each neuron carries about the signal (i.e., song), the generative model (i.e., the prediction of future song), and the prediction-related error. We find strong evidence that the temporal pattern of song-evoked spiking throughout multiple auditory forebrain regions carries information about the predictive features of conspecific song and the error relative to these predictions. In contrast to common computational models of predictive coding (e.g., [1, 12, 4]), neither predictive nor error encoding is segregated in separate neurons. Instead, almost all neurons carry a non-random mixture of both predictive and error information.

## Results

### Natural stimuli as a generative model

Predictive coding theories hypothesize that the brain uses a generative model that maps from a priori causes to predict incoming sensory information. To be minimally functional, this generative model should reflect statistics of behaviorally relevant signals that the animal is most likely to encounter. For songbirds, one class of such signals are the songs of conspecifics. Therefore, as a proxy to a default generative model we modeled the predictive statistics of conspecific song repertoires. To do this, we trained a deep convolutional network that takes starling song as input and learns to predict upcoming starling song acoustics based on recent singing (Figure 1). Following the predictive coding framework [1], we use this generative model to define three representations of the stimulus corresponding to signal, prediction, and prediction-error.

**Figure 1:**
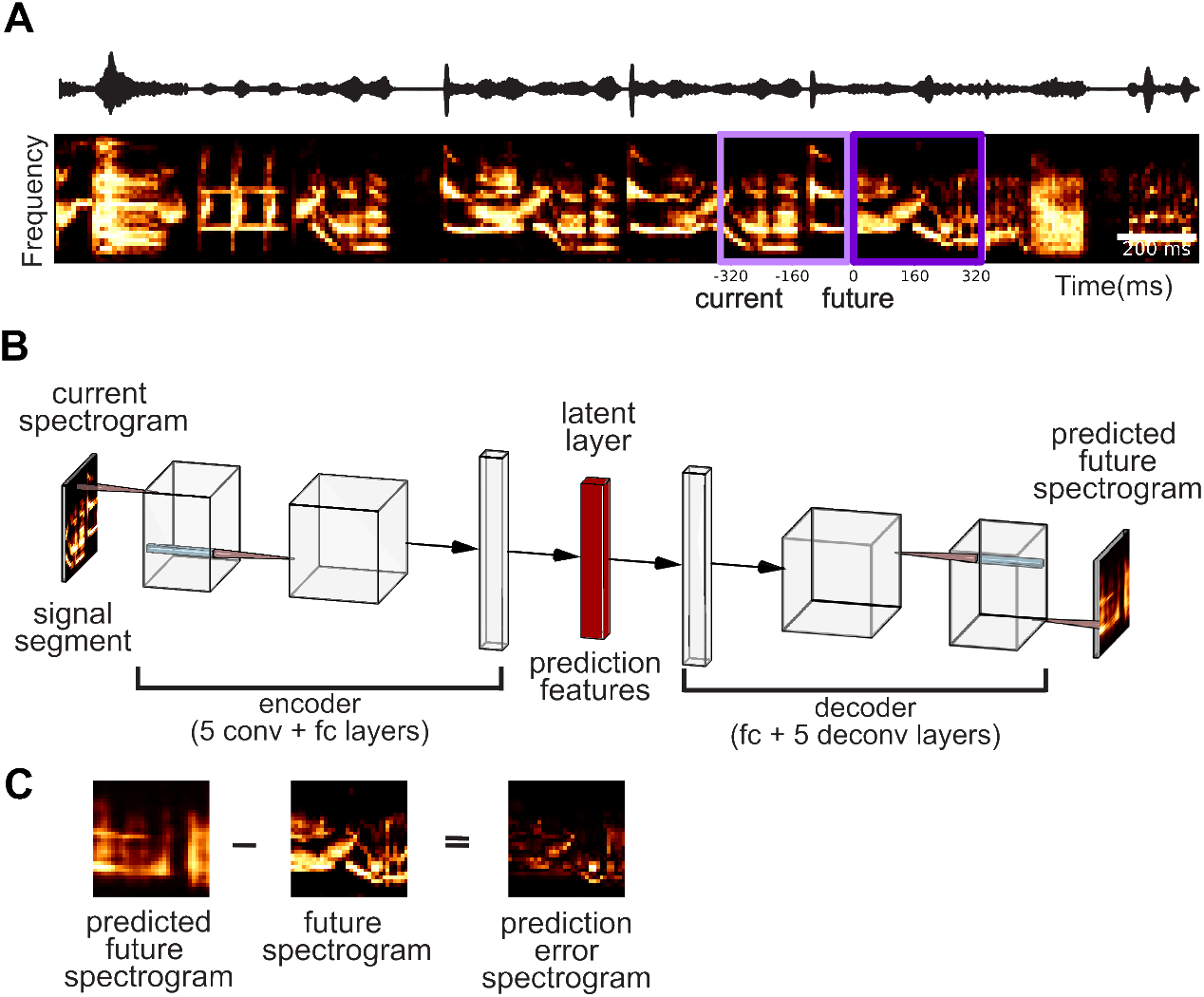
A generative model of the predictive acoustic feature space of European starling song. (A) Example starling song segment showing the time-varying pressure waveform (top) and spectrogram (bottom; frequency range=0.3-12kHz, scale bar=200ms) divided into contiguous “current” and “future” segments each 336msec long (colored squares; 32 frequency channels x 32 time bins). (B) Schematic of the Temporal Convolutional Model (TCM) comprising convolutional and fully connected layers trained on the starling song library to predict future segments (outputs) from current segments of song (inputs). Spectrogram-driven activation of the trained latent layer (256 hidden units) is taken as the prediction representation of the stimulus. (C) Example prediction-error representation taken as the squared difference between true future and TCM-predicted future spectrogram segment.

The network, which we refer to as the Temporal Convolutional Model (TCM), uses a classic encoder/decoder architecture [29] Table S1. It takes a “current” spectrographic segment (336-msec long, 32 frequency bins x 32 time bins) as input, compressing it via the convolutional and fully connected layers of the encoder into a 256-dimensional latent representation, which then passes through the decoder to predict the immediate upcoming, “future”, 336-msec song segment (Figure 1A). Using standard methods to avoid over-fitting (Methods), we trained the TCM to minimize error between predicted and ground-truth future spectrogram segments using a large vocalization library containing over 30 hours of male and female starling song (see Methods, [30], Figure S1A).

We used the trained TCM network to construct signal, prediction, and prediction-error representations for a separate library of natural starling songs that we used for physiological testing. For the signal representation, we simply took the spectrograms of the physiological test songs. To construct prediction representations, we parsed the spectrograms generated from the physiological test stimuli into 336-msec segments and fed them as inputs to the trained TCM network (Methods). This yielded a 256-dimensional latent representation of each input, and a predicted “future” spectrogram based on each input. We used the TCM network latent representation of each input as the prediction representation, with the reasoning that it can be understood as a non-linear transformation of the spectrotemporal song features that are predictive of the upcoming song. We defined the prediction-error representation as the squared difference between the output of the TCM (i.e., the predicted future song segment) and the ground-truth future song segment (Figure 1C). Figure S1B shows examples of these three stimulus representations.

### Mapping auditory responses to signal, prediction, and error representations

According to the strongest form of predictive coding, single neurons should encode either the signal, the prediction from the generative model, or the error that results from comparisons between the signal and prediction [31, 32, 33]. To test this, we recorded conspecific song-evoked spiking activity in anesthetized European starlings from populations of well-isolated single neurons (n=1749 neurons, 8 birds, 14 penetration sites; see Methods, Figure S2) in the primary auditory forebrain region Field L (*n* = 622 neurons, 3 birds), and two secondary auditory forebrain regions, the caudal mesopallium CMM (*n* = 559 neurons, 4 birds) and caudal medial nidopallium NCM (*n* = 568 neurons 3 birds; Table S2). Consistent with prior work [34, 35, 36, 37, 38, 39, 40, 41, 42], robust spiking responses from neurons in all three regions was evoked by presentation of conspecific songs. NCM and CMM neurons displayed precise, sparse activity time locked to specific acoustic features of conspecific song, whereas most Field L neurons responded less selectively to song playback. Figure 2A-C shows example song-aligned spike trains for individual neurons and populations of units in NCM (150 neurons), CMM (*n* = 79) and Field L (*n* = 43).

**Figure 2:**
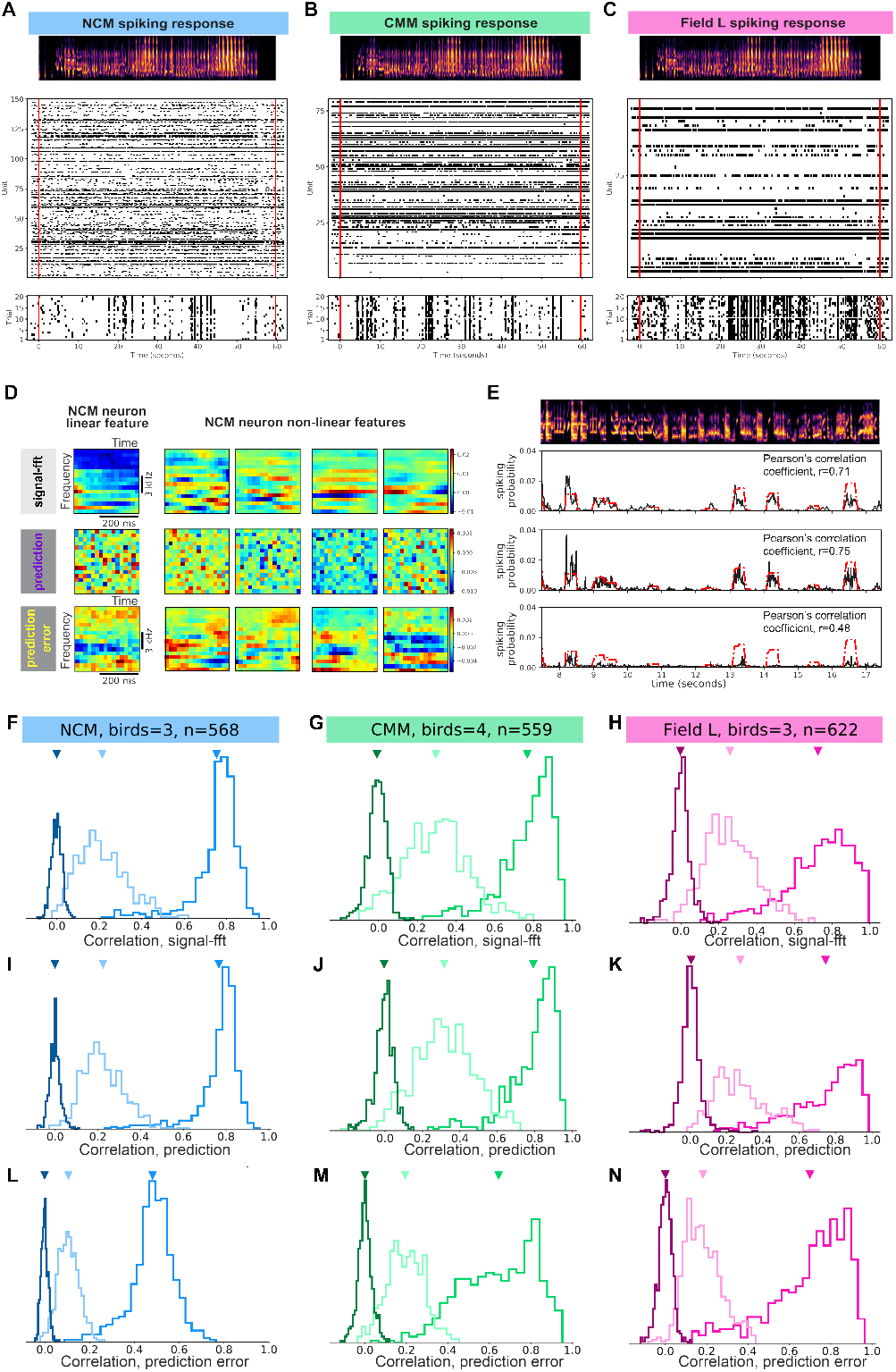
Composite receptive fields of auditory neurons fit to different stimulus representations. Examples of song-aligned extracellular spike trains for single neuron populations in (A) NCM (*n* = 150 neurons), (B) CMM (*n* = 79), and (C) Field L (*n* = 43). Each panel (A-C) shows a spectrogram of an example stimulus (top), neural population spike raster (middle), and a single neuron spike raster (bottom) showing the sequential responses to 20 repetitions of the example stimulus. Red horizontal lines denote the start and end of the stimulus. (D) Example composite receptive fields (CRF; columns) of one NCM neuron fit to three different stimulus representations (rows) showing linear component (left-most column) obtained from parameter h of the MNE model, and four most significant non-linear components (rightmost columns) obtained from MNE parameter J. Features of the prediction-CRF correspond to TCM latent weights, and thus are not acoustically intuitive. (E) Time-varying CRF response predictions for an example novel stimulus (top); CRF-predicted response (black) overlaid on the empirical neuron response (red), with the corresponding Pearson’s correlation coefficient indicating the quality of each CRF model. (F-N) Population-level distributions of Pearson’s correlation coefficients between CRF-predicted and empirical responses for full CRF models (medium saturation), the linear-only component of the model (light saturation), and shuffled control models (Methods; dark saturation). Columns (left to right) show the data for NCM (*n* = 568 neurons from 3 birds), CMM (*n* = 559, 4 birds), and Field L (*n* = 622, 3 birds), respectively. Rows (top to bottom) show distributions generated from the signal-CRFs, prediction-CRFs, and prediction-error-CRFs, respectively. Arrows indicate the position of means in each distribution. The shuffled control models perform significantly worse than all other models (Linear Mixed Effects, see text). Correlation coefficients generated by the various models (signal, prediction, or error) varied significantly across brain regions (see text).

To relate the signal, prediction, and prediction-error representations to neural activity, we computed composite receptive fields (CRFs) using the Maximum Noise Entropy (MNE) method [43, 37, 44], which estimates the probability of a spike given the stimulus. The stimulus is normally taken to be the spectrogram of the response-evoking signal, but in principle may take any representational form, including those that we have defined as the prediction and prediction-error representations. For each neuron, we independently computed the MNE model parameters that optimally relate the neuron’s response to each of the three different stimulus representations: (Methods). We balanced the dimensionality of each stimulus representation and used 80% of the physiological test stimuli for MNE fitting. This yielded a version of each neuron’s CRF fit to either: 1) all spectrotemporal features of conspecific song (signal-CRF) or 2) the predictive TCM latent features of song (prediction-CRF) or 3) spectrotemporal features of prediction-error (error-CRF). The MNE model parameters *h* and *J* correspond to the spike-triggered average and spike-triggered covariance, respectively, giving us linear and non-linear components combined into each CRF. In all cases, the eigenspectrum of *J* contained large, statistically significant eigenvalues, indicating that multiple covariant features of each representation drive spiking responses. Example MNE CRFs generated from the signal, prediction, and error representations for a single NCM neuron are shown in Figure 2D. Because the “receptive field” is a function of the stimulus representation, components of the signal-CRF and the prediction-error-CRF can be displayed in a familiar spectrotemporal coordinate frame. Features of the prediction-CRF are a function of the TCM network latent weights, however, and thus are not readily decipherable as spectrotemporal features. Visual inspection of the CRFs revealed multiple, distinct receptive-field features in individual high-level auditory neurons tied to all three stimulus representations.

The MNE model parameters fit to a neuron’s response can be used to predict the spiking response to novel stimuli. This provides a quantitative estimate of model quality, with better models yielding more accurate response predictions. We predicted the responses of that neuron to the held-out portion (20%) of physiological test stimuli not used for MNE fitting (Methods). We evaluated prediction accuracy for each CRF by computing the correlation coefficient between the modeled response obtained from each stimulus representation and the empirical response of that neuron (Figure 2E). To generate a null distribution of CRF predictions, we repeated the MNE fitting and response prediction procedures using randomly shuffled versions of the spike trains for each neuron. Figure 2F-N shows the distributions of correlation coefficients associated with the CRF response predictions for full (linear/non-linear) models, linear only models, and shuffle controls in each region. The mean correlation coefficient across all the full CRF models (*r* = 0.73 ± 0.004, range over all three regions: 0.70 − 0.75) was significantly higher than that for the shuffled controls (*r* = −5.73*e* − 4 ± 4.98*e* − 4, range: −4.84*e* − 3*to*1.54*e* − 6; Linear Mixed Effects, relative to null distribution across NCM, CMM and Field L: ***β*** = −0.64, *SE* = 0.008, *z*(7970) = −77.210, *p* < 2*e* − 16; Figure 2F-N). The same was true for specific CRF models within each region: NCM signal-, prediction-, and error-CRFs, *t* > 116.2, *p* = 0.0; CMM, *t* > 83.9, *p* = 0.0; Field L, *t* > 91.7, *p* = 0.0, paired t-test all cases relative to corresponding null distributions; Figure 2F-N). In summary, all the composite receptive fields, regardless of the stimulus representation with which they are fit, model portions of the empirical responses in NCM, CMM and Field L better than chance.

### Predictive and prediction-error responses vary across regions

The MNE receptive fields computed here allow for quantitative comparison of how distinct predictive and prediction-error features of song are tied to forebrain responses. The pattern of correlation coefficients produced by the different models (signal, prediction, or error) varied significantly across brain region (NCM, CMM or Field L; Linear Mixed Effects, NCM vs CMM: ***β*** = −0.263, *SE* = 0.036, *z*(3990) = −7.296, *p* < 1*e* − 4; NCM vs Field L: ***β*** = −0.219, *SE* = 0.038, *z*(3990) = −5.816, *p* < 1*e* − 4; Field L vs CMM: ***β*** = −0.044, *SE* = 0.033, *z*(3990) = −1.312, *p* = 0.388). Across all regions, both the signal- and the prediction-CRFs yielded excellent models for the empirical responses of most neurons to conspecific song, accounting for an average of 55.87% and 58.80% of the response variance, respectively. By comparison, while still well above chance, the error-CRFs yielded poorer models, accounting for an average of 37.30% of the total response variance. Within each region the prediction-CRFs yielded the best stimulus-response models, performing slightly but significantly better than the signal-CRFs in all cases (NCM: *t* = 7.7, *p* = 6.76*e* − 14; CMM: *t* = 12.1, *p* < 1*e* − 23; Field L: *t* = 11.2, *p* < 1*e* − 23; paired t-test vs. signal-CRF; Figure 3). In general, both the signal- and prediction-CRFs performed better than the error-CRFs. Correlation coefficients for the error-CRFs were lowest in NCM (mean *r* = 0.48 ± 0.004), and significantly higher by comparison in both CMM (mean *r* = 0.64 ± 0.008; *t* = −19.1, *p* < 1*e* − 23, t-test CMM vs NCM) and in Field L (mean *r* = 0.70 ± 0.008; *t* = −24.9, *p* < 1*e* − 23, t-test Field L vs NCM; Figure 3). In Field L, the response predictions for the error-CRFs were nearly as strong as those for the prediction- and signal-CRFs. Not surprisingly, combining the signal-, prediction- and error-CRFs yields a prediction of song-evoked responses that is better than any single model (NCM, 67.31 ± 0.004% explained variance, *t* = 55.36, *p* = 7.6*e* − 231; CMM, 67.93 ± 0.004% explained variance, *t* = 58.94, *p* = 3.87*e* − 238; Field L, 67.88 ± 0.04% explained variance, *t* = 63.00, *p* = 2.02*e* − 271 paired t-test vs best single model). This supports the hypothesis that neurons in all three forebrain regions carry information about predictive features of the signal along with the signal-driven error relative to a model of conspecific song, with the strength of the prediction-error representation diminishing between Field L and NCM.

**Figure 3:**
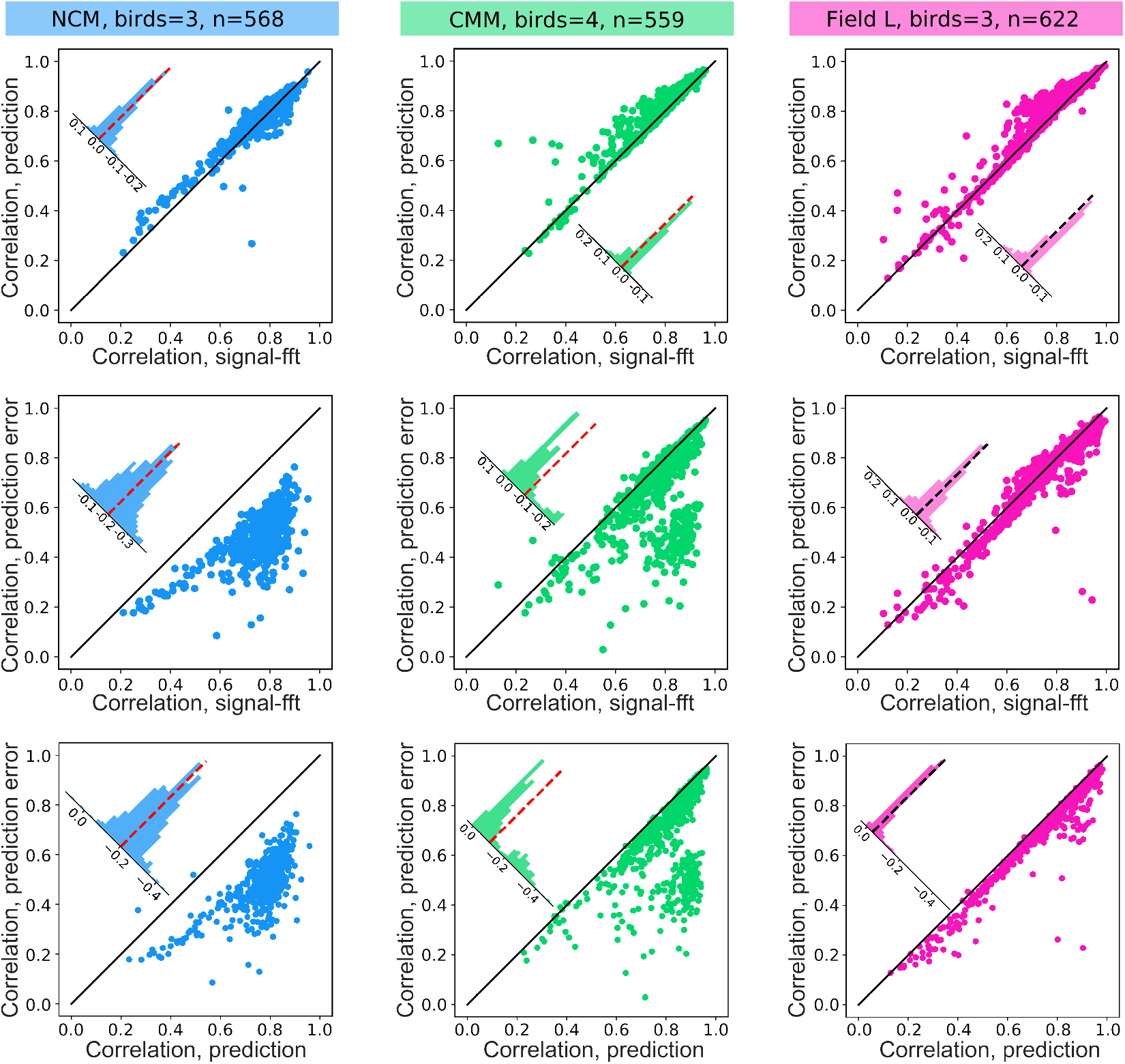
Relative strength of encoding for signal, prediction, and prediction-error features in single auditory neuron responses. Scatterplots showing comparative CRF-model quality (Pearson’s correlation coefficient values between CRF-predicted and empirical responses to test stimuli) in single neurons. Columns (left to right) show the data for NCM (*n* = 568 neurons), CMM (*n* = 559), and Field L (*n* = 622), respectively. Rows (top to bottom) show comparisons between prediction-CRFs and signal-CRFs, prediction-error-CRFs and signal-CRFs, and prediction-CRFs and prediction-error-CRFs, respectively. The black diagonal line indicates unity. The distribution of residuals is inset in each plot with a dashed line indicating the mean residual.

We note in addition that the relative strength of this song prediction-error processing in CMM and Field L is closely tied to the non-linear component of the CRFs in these regions. Not surprisingly, the full MNE models enable significantly better response predictions than those produced from only the linear component of the CRFs (paired t-tests, *p* < 1*e* − 23 all cases; Figure 2F-N). Within the context of this main effect, however, we observed a significant interaction between stimulus representations (signal, prediction and prediction-error), MNE models (linear or full model) and brain region (NCM, CMM and Field L; 3-way interaction terms, *F* (4) = 60.018, *p* < 1*e* − 4), that is driven primarily by the fact that the non-linear CRF features add much less error information in NCM compared to CMM and Field L (Figure S3).

### Alternate generative models

To validate performance of the prediction-CRFs which are fit to a latent stimulus representation, we trained an autoencoder network [28, 45] that shares a similar architecture with the TCM but reconstructs the current spectrographic input segment rather than the future segment (Methods). Like the TCM, the autoencoder latent space is a non-linear compression of spectrotemporal song features, but presumably contains features that are both predictive and non-predictive of upcoming song, as does the spectrogram itself. Like the prediction-CRF, the autoencoder-CRF (ae-CRF) does not display intuitive spectrotemporal structure. Like the prediction-CRFs, the ae-CRFs yield very high-quality models of the empirical spiking response, similar to that achieved with the spectrographic representation (Figure S4). Within each region the ae-CRFs performed well above chance (*t* = 162.8 NCM; *t* = 130.6 CMM; *t* = 94.1 Field L; *p* = 0.0 all cases, Figure S4J-L). Across all three regions, the mean Pearson’s correlation coefficient between the empirical and ae-CRF predicted response was *r* = 0.79 ± 0.005, similar to the mean across all three regions for the signal-CRFs (*r* = 0.75 ± 0.005). Thus, the latent space of networks such as TCM can provide valid stimulus representations. In practice, because forebrain auditory regions are biased to song [40], projecting song into any network latent space should yield CRF models that are better than chance, so long as the network transformation of song is repeatable (i.e. the same song input yields the same latent representation). If neurons throughout the songbird auditory forebrain are optimized for predictive features of conspecific song, however, then the latent space of the song-trained TCM network should yield better models than the same network trained on other (non-song) inputs. To test this hypothesis, we repeated our CRF fitting procedure using the TCM network trained on MNIST digit sequences rather than starling song (Methods). This yielded prediction-CRFs that were significantly poorer models of the empirical spiking responses to song compared to those for the song-trained TCM (*p* < 1.1*e* − 62, *t* > −18.8, all cases, paired t-test; Table S3), consistent with the idea that neurons are preferentially tuned to predictive features of song. We also tested different network architectures as proxies for the internal generative model. Both a simple feedforward neural network (Temporal Predictive Model or TPM) and a variation of the Temporal Convolutional Model (TCMv2; see Methods; Figure S5A-C) yielded similar results as those reported above. Like TCM, the TCMv2 architecture generated higher Pearson’s correlation coefficients for modeled responses in comparison to signal-CRFs (NCM, *n* = 214; Linear Mixed Effects, contrasts between architectures, TCMv2 vs signal: ***β*** = 0.03, *SE* = 0.005, *z*(3850) = 6.171, *p* < 1*e* − 06; TPM vs signal: ***β*** = −0.0004, *SE* = 0.005, *z*(3850) = −0.105, *p* = 0.994; Figure S5D).

### Shared and unique prediction and prediction-error encoding

The foregoing analyses support the idea that song-evoked responses in NCM, CMM, and Field L carry information about the signal, predictions, and prediction-errors. All three representations are a function of the same stimulus, however, and it is therefore likely that a portion of the information captured by each is redundant. To the extent that the different CRFs describe non-independent components of the stimulus-response relationship, their labelling is arbitrary. To investigate this, we partitioned the variance of the stimulus response predicted by each CRF model, separating the explained response variance into unique portions attributable to the signal-, prediction-, and error-CRFs, along with shared portions common to pairs of, or all three, CRFs. We used a regression model for the partitioning strategy, treating the empirical neural response as a composition of modeled response probabilities (Methods). Figure 4 shows the average shared and unique variance partitions in each of the three forebrain auditory regions. Importantly, all the partial correlation coefficients were significantly greater than that expected by chance (NCM, *p* < 0.05; CMM, *p* < 0.05; Field L, *p* < 0.05; two-sample Kolmogorov-Smirnov test; Table S4). Somewhat surprisingly, however, given the nature of each representation, the proportion of response variance shared between all three CRFs was relatively low overall (NCM, *partialr* = 0.04 ± 0.001; CMM, *partialr* = 0.12 ± 0.005; Field L, *partialr* = 0.18 ± 0.004), and significantly lower in NCM compared to CMM (Wilcoxon rank-sum test, *p* = 3.74*e* − 66, Cliff’s *d* = −0.595) and Field L (Wilcoxon rank-sum test, *p* = 1.91*e* − 175, Cliff’s *d* = −0.947; Table S4, Figure S6). The single overall largest partition in each region was that shared by the prediction- and the signal-CRFs (NCM, *partialr* = 0.51 ± 0.004; CMM, *partialr* = 0.63 ± 0.008; Field L, *partialr* = 0.66 ± 0.007; Table S4, Figure S6). Like the full correlations, within each region the single largest unique variance partition was attributable to the prediction-CRF (NCM partial *r* = 0.39 ± 0.004; CMM partial *r* = 0.29 ± 0.008; Field L *partialr* = 0.23 ± 0.007), which was significantly larger than the unique partition for the signal-CRF (NCM, *t* = 11.3, *p* < 1*e* − 23; CMM, *t* = 10.5, *p* = 1.2*e* − 23; Field L, *t* = 15.6, *p* < 1*e* − 23 paired t-tests) and the error-CRF (NCM, *t* = 72.2, *p* < 1*e* − 23; CMM, *t* = 19.0, *p* < 1*e* − 23; Field L, *t* = 23.9, *p* < 1*e* − 23 paired t-tests; Figure 4).

**Figure 4:**
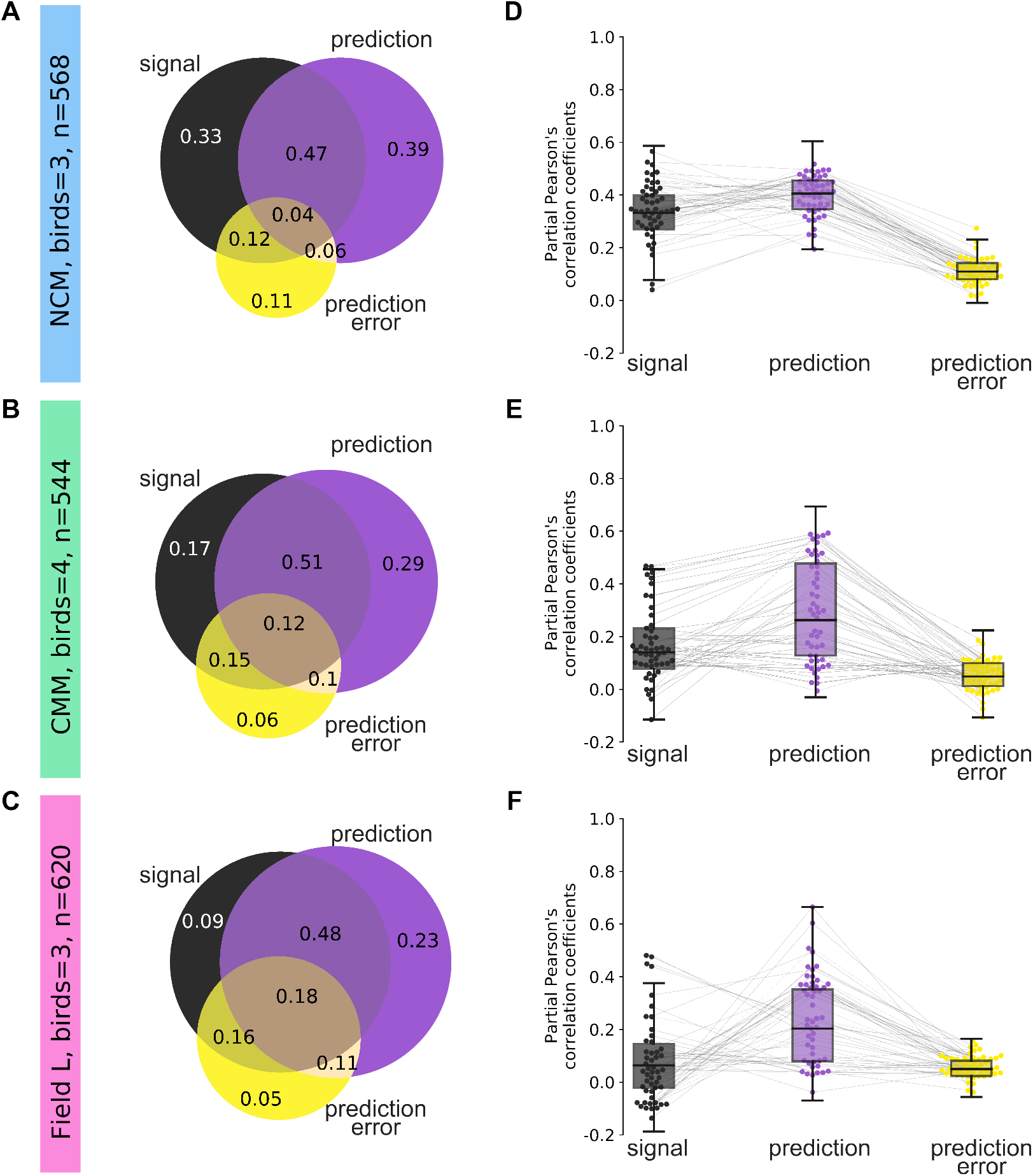
Unique contributions of signal, prediction and prediction-error varies across auditory regions. (A-C) Visualization of the mean unique and shared variance (partial Pearson’s correlation coefficients) attributable to the signal (black), prediction (purple) and prediction-error (yellow) representations of song in NCM (n=568 neurons), CMM (n=544) and Field L (n=620). (D-F) Boxplots showing overall distribution of mean partial correlation coefficients uniquely attributable to the signal, prediction and prediction-error representation in NCM, CMM, and Field L. Overlaid swarm plots show a random sample of the same 50 neurons in each region. The highest unique response prediction representation is associated with the highest unique response variance in each of the three regions (see text).

Partitioning of the response variance revealed prediction-error differences across the three brain regions. The pattern of unique response variance associated with each stimulus representation varied significantly between regions (Two-way analysis of variance (ANOVA), main effect for stimulus representation, *F* (2, 5187) = 1522.68, *p* < 2.2*e* − 16, main effect for brain region, *F* (2, 5187) = 743.98, *p* < 2.2*e* − 16, stimulus representation × brain region interaction term, *F* (4, 5187) = 88.14, *p* < 2.2*e* − 16). In NCM the total response variance attributed to prediction-error was significantly smaller (*r* = 0.48 ± 0.004) compared to CMM (mean *r* = 0.64 ± 0.008) and Field L (mean *r* = 0.70 ± 0.008; *p* < 1*e* − 23 both cases, t-test; Figure 3). At the same time, however, the unique error partition (i.e. the response variance attributed only to the error representation) was significantly larger in NCM (partial r=0.11) compared to CMM (partial *r* = 0.06, *t* = 15.1, *p* = 6.2*e* − 47) and Field L (partial *r* = 0.05, *t* = 20.4, *p* = 2.8*e* − 7, t-test between regions; Figure 4, Table S4, Figure S6). This suggests a general processing hierarchy between Field L/CMM and NCM across which the overall strength of error encoding decreases, while predictive and prediction-error components become increasingly dissociable in the neural response.

### Prediction and prediction-error encoding is mixed in single neurons

In all three regions, significant components of the song-evoked response can be uniquely attributed to acoustic features of the signal, prediction, and prediction-error representations. Many computational theories of predictive coding posit that these different responses will be segregated to functionally specialized cell-types, e.g., “error coding” neurons [31, 4, 46, 47, 48]. If such specialization is a general property of neurons within a brain region, then a negative relationship should exist between the unique partial correlation coefficients attributed to prediction and prediction-error driven responses of each neuron. Such relationships are, however, not prominent in our samples. On average in each region, the relationship between prediction and error encoding is either flat or slightly positive (NCM, Spearman’s *r* = 0.23, *p* = 1.94*e* − 8; CMM, Spearman’s *r* = 0.09, *p* = 4.17*e* − 2; Field L, Spearman’s *r* = 0.33, *p* = 2.79*e* − 17; Figure 5A), contradicting the region-wide functional specialization hypothesis. Nonetheless, the presence of significant correlations in each region indicates that predictive and error encoding strength in single neurons within each region are neither independent nor random. To examine functional specialization in single neurons further, we took the difference between the partial correlation coefficients associated uniquely with the error and predictive representations as a simple measure of encoding bias in each neuron. As expected, given the relative strength of the mean prediction compared to error encoding, the mean residual (error minus prediction) in each region was negative (NCM=−0.28 ± 0.004; CMM=−0.24±0.008; Field L=−0.18±0.006; Figure 5B). In the limit, functional specialization for either the prediction or error implies that the bias residual for a neuron should lie in either the negative or positive tail of the overall residual distribution, respectively. We observe little support for the possibility of exclusive error processing, as the positive tails in NCM and CMM (relative to the 95% CI) are not significantly larger than expected by chance (*X*2(1), *p* > 0.26, both cases) and in Field L the positive tail is significantly smaller than expected by chance (*X*2(1) = 0.002, *p* = 0.03; Figure S7; Table S5). In terms of exclusive predictive processing, the negative tails of the residual distributions (relative to the 95% CI) for NCM and CMM again do not differ from chance (*X*2(1), *p* > 0.06, both cases). In Field L, however, we observe significantly more neurons in the negative tail than expected by chance (*X*2(1) = 0.003, *p* = 0.04), consistent with a small population (ca. 5% of neurons) that may be functionally specialized for predictive coding (Table S5).

**Figure 5:**
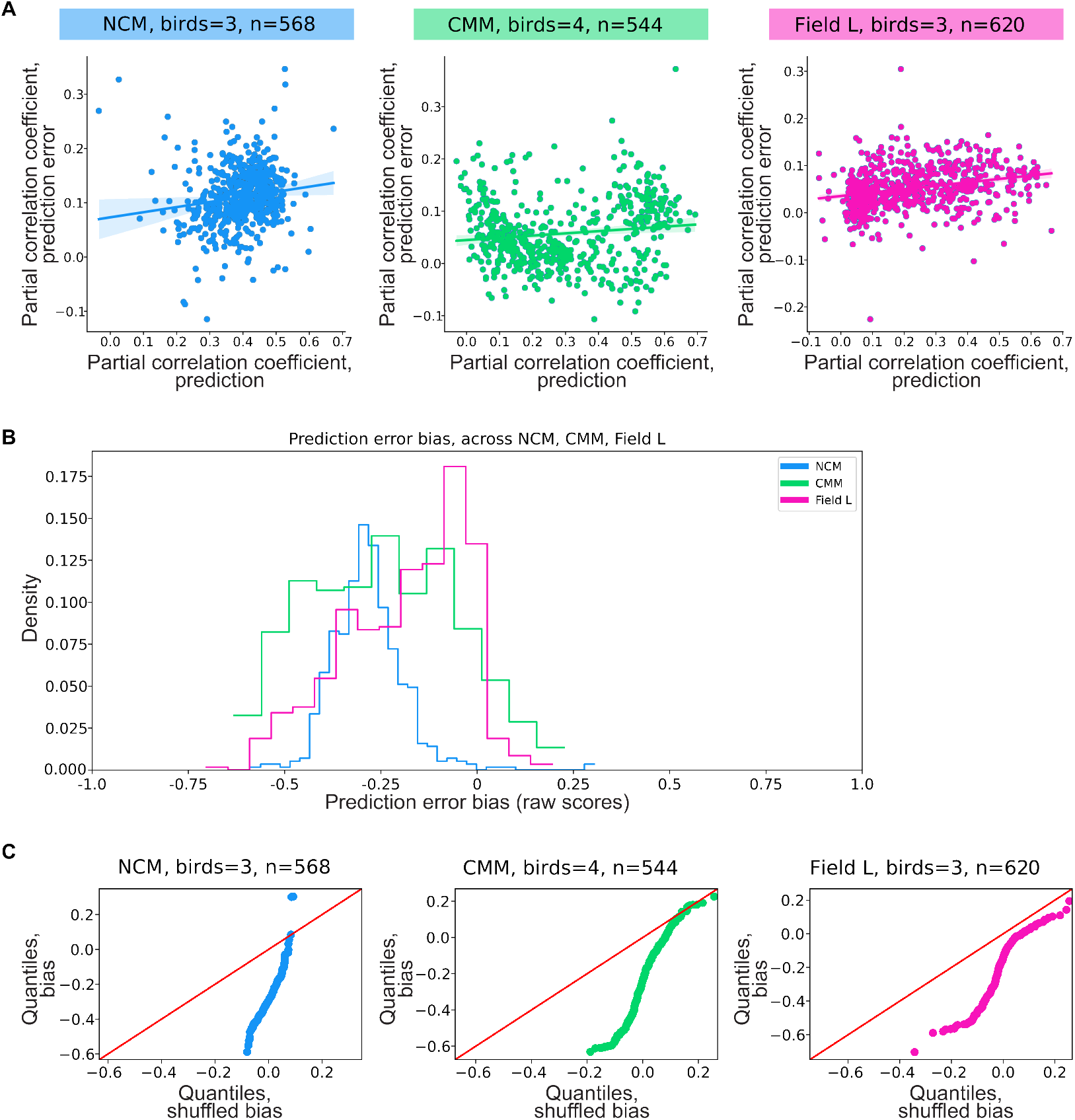
Prediction and error bias in single neurons. A) Scatterplots showing relationship between prediction and error encoding in NCM, CMM and Field L single neurons. In each region, the relationship is either flat or slightly positive (see Text for stats). (B) Histograms showing distribution of bias residuals, determined by the difference between unique error and unique prediction (error - prediction), in NCM, CMM and Field L. The residual distributions differ significantly between regions (ANOVA, F=61.42, p=1.81e-26). (C) Quantile-quantile (q-q) plots comparing bias residual distributions from empirical and shuffled CRFs in each neuron. Each point shows the values for the corresponding quantile in each distribution, red lines indicate unity, and non-linearities show deviations.

In contrast to the small numbers of functionally specialized neurons in each region, most neurons carry information about a combination of predictive and error components of song. To understand the relationship between predictive and error information within each neuron, we created quantile-quantile (Q-Q) plots of the bias residuals in each region (Figures 5, S7). These plots capture deviations from the corresponding shuffled bias scores, but across the full distributions. Overall, the empirical bias distributions deviate significantly from normal (*p* < 1.05*e* − 5, all cases, Shapiro-Wilk test; Table S6), and the nonlinear structure of the Q-Q plots (Figure 5C) indicate that none of the distributions is a simple linear translation of the shuffle control. Finally, we note that the bias residual distributions differ significantly between regions (ANOVA, *F* = 61.42, *p* = 1.81*e* − 26, Figure 5). Thus, the combination of predictive and error information in individual neurons in each region is not random, and the specific non-random mixture varies between regions. To examine the possibility that more complex, but still dissociable, response profiles might exist in our sampled populations, we applied a hierarchical density-based clustering algorithm (HDBSCAN; [49]) to the unique partial correlations attributable to signal, prediction and prediction-error features and quantified cluster strength in each region. Figure 6 shows example clusters for single simultaneously recorded populations in each of the three regions, along with mean silhouette scores for our full dataset. Across the entire sample populations, we observed more distinct clustering for predictive and non-predictive features in both CMM and Field L, compared to NCM. Clusterability was strongest in Field L (mean silhouette score= 0.47) and CMM (mean= 0.36), and lowest in NCM (mean= 0.20), and varied significantly between regions, (ANOVA, *F* = 5.33, *p* = 0.02; Figures 6, S8).

**Figure 6:**
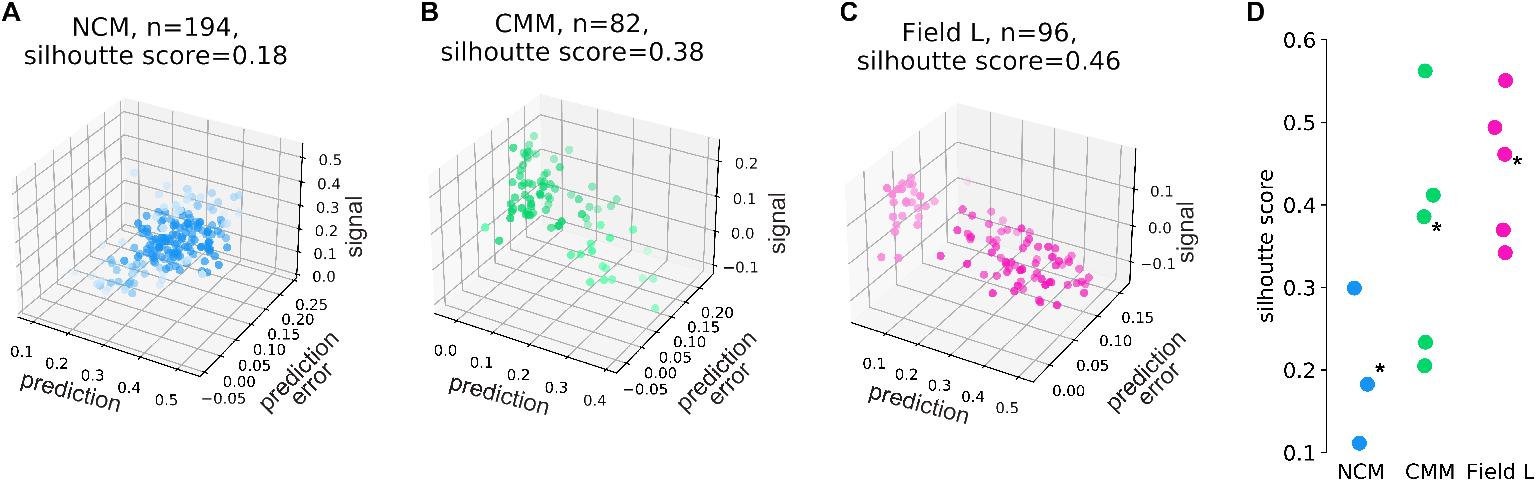
Response clustering based on signal, prediction and prediction-error driven responses. Three-dimensional scatterplots showing unique response variance (partial Pearson’s correlation coefficients) attributable to signal, prediction, and prediction-error representations in example populations of simultaneously recorded neurons in (A) NCM (n=194 neurons) (B) CMM (n=82) and (C) Field L (n=96). (D) Silhouette scores based on HDBSCAN clustering (Methods) for all recorded populations in NCM (n=3), CMM (n=5) and Field L (n=6). Higher silhouette scores indicate stronger clusterability. Asterisks mark the populations shown in A-C. Clusterability was strongest in Field L and varied significantly between regions (see text).

## Discussion

We combine computational modeling with electrophysiology to understand the neural representation of vocal communication signals in the songbird auditory forebrain from the perspective of predictive coding. Using the natural statistical structure of conspecific song as a proxy for an internal generative model, we show that spiking responses of single neurons throughout primary and secondary auditory forebrain regions are reliably driven by the predictive features of conspecific song and the error in explicit vocalizations relative to these predictions continuously in time. Collectively, our results bridge the gap between qualitative evidence supporting different predictive coding theories and more mechanistic understanding of how such theories are implemented. Conceptual theories of predictive coding generally follow one of two forms. The first, which derives from theories of efficient coding and has empirical support in multiple sensory cortices, focuses on prediction error [50, 51, 52, 53, 54, 55]. In the auditory system, responses to deviant, omitted, or surprising stimuli, and adaptation to repetitive sounds [56, 57, 58, 59, 60, 61, 17] is used to support the notion that neurons encode a prediction error [18, 19, 20] derived from the difference between the current input and an internally generated expectation. The expectation may be a product of the stimulus history (e.g. a repeated tone) or other learned contingencies [22, 23]. A second, non-equivalent form of predictive coding, examined primarily in retinal visual processing [62, 63], posits that sensory systems are adapted to preferentially encode sensory information that is predictive of the future [64, 63]. These two forms of predictive coding are distinguished by the information carried in the neural responses. Error driven responses, of the kind evoked by surprising or deviant stimuli are informative about the past, whereas predictive responses are informative about the future [13]. Although distinct, these two notions of predictive coding are not incompatible, and both forms of information are clearly useful. Our results support both notions of predictive coding in the context of a complex natural vocal communication signal, showing that responses of single neurons carry information about the predictive features of the vocal signal and their error relative to species-specific model.

Predictive coding has been proposed as a general cortical mechanism [3, 4] including for speech processing [5, 6]. To function in this context, it must implement continuous, real-time predictions and errors. Our use of a neural network as a proxy for a species-specific internal model was crucial to building a continuous predictive representation of song. On average, our predictive networks outperform classical spectrographic representations (Figures 3, 4). Prior work has argued that similar predictive networks, trained on natural stimuli, can account for basic features of receptive fields in auditory and visual cortices [65, 61]. The high quality of the receptive field models fit directly to the latent features of the predictive network here, provide direct empirical confirmation for the utility of these networks in revealing useful stimulus representations. Although we experimented with several network architectures that yielded similar results (Figure S5) and show that song-trained networks outperform those trained on arbitrary sequences, we did not attempt to find an optimal network. Future work to find more optimal networks could directly exploit models for the sequential acoustic structure of song [11] along with recently developed network approaches that directly incorporate temporal information [66], or those that seek to jointly optimize the fit of responses and the structure of the predictive latent space.

Prior work in songbirds supports error encoding in NCM [67, 57, 54]. Our results confirm this earlier work and extend it by showing that both prediction and error are uniquely encoded by neurons throughout the songbird auditory forebrain. Across forebrain regions, we observe some heterogeneity in the mixture of prediction and error encoding that is consistent with a general processing hierarchy between Field L/CMM and NCM across which the overall strength of error encoding decreases. More obvious, however, is the fact that nearly all single neurons in these regions carry both predictive and error information. This observation contradicts the standard formulation for models of the predictive coding, which posit separate prediction and prediction-error encoding neurons [4, 32, 31]. More recent theoretical work shows, however, that error computations can be performed in dendritic compartments [68, 69, 70, 71] rather than specialized neurons, and that predictive and error responses can emerge spontaneously in the same neurons in recurrent neural networks [72]. Rather than thinking of neurons as carrying a “representation” of either the predictive features or the error, we prefer to conceive of the spiking responses as effectors that drive the system toward behavioral goals. That is, the utility of the error likely serves a behavioral rather than representational goal. Testing this idea requires measurement of prediction and error responses while the animal weighs competing internal models tied to different behavioral goals.

## Acknowledgements

We thank Marvin Thielk, Tim Sainburg for critical discussions and/or input on the manuscript. This work was supported by the US National Institutes of Health grant R01DC018055 to T.Q.G. Conceptualization, S.R. and T.Q.G.; methodology, S.R. and T.Q.G.; investigation, S.R., M.E.T. and B.H.T.; software, S.R. and B.H.T.; formal analysis, S.R.; data curation, S.R.; visualization, S.R. and T.Q.G.; writing: original draft, S.R. and T.Q.G.; writing: review & editing, S.R. and T.Q.G.; resources, T.Q.G.; supervision, T.Q.G.; funding acquisition, T.Q.G. The authors have no competing interests to declare.

## Methods

### Animal subjects

Under a protocol approved by the Institutional Animal Care and Use Committee of the University of California, San Diego, we collected electrophysiology data from n=8 adult (>1 yrs. old) European starlings (*Sturnus vulgaris*). All birds were wild caught in southern California and maintained prior to physiological testing on a light/dark cycle matched to the ambient photoperiod. Food and water were freely available. We did not control for the sex of the subjects.

### Physiological test stimuli

Starlings (both male and female) produce long, spectrotemporally rich, individualized song bouts comprising repeating, shorter acoustic segments known as “motifs” that are learned over the bird’s lifespan. Motifs are on the order of 0.5–1.5 s in duration, and song bouts can many 10’s of seconds. During electrophysiological recording sessions, we presented five one-minute-long natural song bouts and multiple short synthetic ‘songs’ consisting of 1-5 natural starling motifs. The songs and motifs were manually extracted from a larger library of natural starling songs produced by several singers. The synthetic songs were created using motifs from eleven singers. The singers featured in our test stimuli are different from the ones employed for neural network training. Original song samples were recorded from European starlings at either 44.1 or 48kHz (16-bit).

### Electrophysiology

Experimentally naive starlings were anesthetized with urethane (0.7 mg/kg) and head-fixed to a stereotaxic apparatus in an acoustic isolation chamber (Acoustic Systems). A small craniotomy was opened over the target region NCM, CMM or Field L. We placed multi-channel silicon electrodes (Masmanidis 128DN; [73]) in the region of interest until auditory-evoked activity was observed on many channels. Songs were played to the subjects at 60-dB SPL mean-level while we recorded action potentials extracellularly. After placing the electrode, the bird was left in silence for 30-60 minutes prior to starting presentation of the test stimuli. Each test stimulus was repeated 20 times in a random order with random inter-trial intervals of 2-5 seconds. When possible, given the stability of the preparation, recording blocks were obtained from independent populations of cells in both hemispheres and at different well-spaced depths within the region of interest (Table S2).

### Temporal Convolutional Model

We trained a neural network model to generate predictions of song on the timescale of the approximate average length of starling song motifs. Specifically, the network takes a 336ms song segment as input and predicts the next immediate future song segment of the same length. We propose a deep convolutional network [45] Temporal Convolutional Model (TCM) trained to predict future segments of song as a proxy to the generative model in the brain (Figure 1B, Table S1). The architecture of this model used a combination of convolutional layers and fully connected layers in the encoder and decoder; and has been shown to resemble neuronal processing in the visual system [74, 75, 29, 76, 77]. The encoder compresses the past spectrographic segment taken as inputs (32×32 units) into a latent representation (256 hidden units), which is then passed through the decoder to predict the output future spectrogram segment (32×32 units). We used a loss function that is a combination of prediction loss and multidimensional scaling (MDS) distance loss. Prediction loss minimizes the mean squared error between the predicted future spectrogram segment and ground-truth future spectrogram segment. The multi-dimensional scaling (MDS) algorithm tries to minimize information loss when projecting data into lower dimensions. It is a non-linear optimization problem which tries to optimize the mapping in the target dimension based on the original pairwise distance information, thus preserving the global structure [78].

### TCM training dataset

We used a large library of previously recorded natural starling songs [30] to create the Temporal Convolutional Model (TCM) network training stimuli. Original song samples in the source library comprise over 30 hours of song with more that 90,000 separate syllables recorded from six European starlings at either 44.1 or 48 kHz (16-bit) in sound-isolated chambers over the course of several days to weeks, at various points throughout the year. Some birds were administered testosterone before audio recordings to increase singing behavior. We down-sampled each song waveform in the source library to 24 kHz and computed its corresponding STFT spectrogram (NFFT=128, a Hanning window of length 128, and 50% window overlap), excluded the DC component, and log-scaled the spectrogram magnitudes. To reduce their dimensionality, we averaged pairwise neighboring time bins twice and frequency bins once as in [37] yielding spectrograms with 32 frequency bins and approximately 10.5 msec time bins. We then parsed spectrograms into segments of 32 time bins in length (ca. 336 msec), with neighboring segments offset by a single time step (10.5 msec). From the resulting set, containing over 10 million 1024-dimensional spectrogram segments, we chose 844399 segments at random to train the TCM network. We paired each input spectrogram segment with the immediately following, non-overlapping segment from the same song, which became the output training target of the TCM network (Figure 1A). We refer to the input and output segments as the “current” and “future” segments, respectively. We reserved 10% of the training set as a validation subset.

### Architecture

The TCM neural network (Figure 1B) comprises both an encoder and a decoder. The encoder is a combination of 5 convolutional layers and a fully connected layer, projected to a 256-dimensional latent space, and a mirror image decoder with a fully connected layer and 5 deconvolutional layers (Table S1). Convolutional layers used different numbers of 3’3 filters and rectified linear unit (ReLU) as the nonlinear activation function. Batch normalization was not used. The TCM network served as our proxy for the internal generative model, and we used the 256-d latent space to represent the features of any given input (current segment) that are most predictive of the output (future segment). As a control representation of all the spectrotemporal features in song, we trained a convolutional autoencoder to the TCM network, except that it learned to reconstruct the current (input) segment rather than the future. As a separate control, we trained the TCM network on digit sequences built from the MNIST dataset (http://yann.lecun.com/exdb/mnist/).

We implemented, trained, and tested the TCM and autoencoder networks described above using Tensorflow [79]. The convolutional and linear weights were initialized to be uniformly random (the default setting in Tensorflow). The loss function for TCM is a sum of a “prediction loss” and “distance loss”. We defined prediction loss as the mean squared error between the true and predicted output, and distance loss using a Multi-dimensional Scaling (MDS) function that maximally preserves the pairwise distances between input and output data points and favors the preservation of global dataset structure. The architectural setting of our autoencoder model was similar to the TCM with stacked convolutional layers. The autoencoder loss function is a sum of “reconstruction loss”, defined as the mean squared error between true and reconstructed output, and MDS distance loss. The models were trained with the ADAM optimizer with a learning rate 0.001 and using mini-batches of size 128 and without using dropout regularization. All additional architectural details can be found in the GitHub repository. The autoencoder followed those in our AVGN repository (https://github.com/timsainb/AVGN).

### Signal, prediction, and prediction-error representations

We computed all 32×32 spectrogram segments from the physiological test stimuli and passed them through the trained TCM to produce a predicted future spectrogram segment. The current spectrogram segment corresponds to signal representation. We used the TCM latent space as predictive representation; and pixel-wise squared difference between the predicted future spectrogram segment and true future spectrogram segment as the prediction-error representation (Figure 1C). Other methods prediction error metrics, e.g., difference error or absolute error, yield similar results (Figure S9).

### Alternative generative models

#### Temporal Prediction Model (TPM)

We implemented (in Tensorflow) a temporal prediction model, inspired by [61], using a feedforward neural network with a single hidden layer of nonlinear units (256 units), designed to map inputs to outputs through weighted connections (Figure S5B). Each hidden unit first performs a linear transformation of past input data (spectrogram segments, 32 frequency bins x 32 time bins) using input weights, followed by a monotonic nonlinearity (sigmoidal used here). The outputs from these hidden units are then linearly mapped to predict future values (spectrogram segments, 32 frequency bins x 1 time bin) using output weights. This approach aligns with the observation that neural decoding can often be approximated by linear transformations. The model architecture is a standard fully connected feedforward network, where each hidden unit computes a weighted sum of its inputs, which is then transformed using a nonlinear activation, such as a sigmoidal function. This setup allows the model to capture both linear and nonlinear relationships in the data for prediction accuracy.

#### Temporal Convolutional Model version 2 (TCMv2)

Another version of the TCM neural network (Figure S5B) comprises both an encoder and a dual decoder. The encoder is a combination of 3 convolutional layers and a fully connected layer, projected to a 256-dimensional latent space, and a mirror image decoder with a fully connected layer and 3 deconvolutional layers to reconstruct the input segment (340 msec of song, spectrogram segments, 32 frequency bins x 32 time bins), and another decoder with only a single fully connected layer in order to predict the future song (about 10.5 msec of song, immediate next spectrogram segment, 32 frequency bins x 1 time bin). Convolutional layers used different numbers of 3’3 filters and rectified linear unit (ReLU) as the nonlinear activation function. The network was implemented in Tensorflow with an ADAM optimizer. The distance loss function, defined using a Multi-dimensional Scaling (MDS) function, was used for decoder to reconstruct input segment, whereas the prediction loss function defined as the mean squared error between the true and predicted output was used for the other decoder. For estimating optimum size of latent space, we tested hidden layers with 1024, 512, 256, 128, 64 and 32 units. We found 256 units to be the most optimum (Figure S5C).

### Electrophysiology data acquisition

Raw extracellular voltages recorded from each microelectrode site were amplified and digitized at 30 kHz through a headstage (RHD 128-Channel Headstage, Intan Technologies) output to an OpenEphys recording system. For online visualization, electrical signals were bandpass filtered between 300 and 6000 Hz and further normalized using common median referencing. We saved the full extracellular voltage waveforms for offline spike-sorting and other analyses along with waveforms of the playback audio (physiological test stimuli) synchronized with the OpenEphys data acquisition streams.

### Spike Sorting

Simultaneously recorded blocks were spike sorted with Kilosort2 [80] followed by manual curation. Automatic sorting was performed in Matlab (Mathworks), and putative units were manually curated using the *phy* interface. Single units were identified by clustering principal components of the spike waveforms, only when fewer than 1% violations of the refractory period (assumed to equal 2 msec) occurred, and only from recordings with an excellent signal-to-noise ratio (large-amplitude extracellular action-potential waveforms). The remaining spike clusters were classified as multi-unit activity or noise based on signal-to-noise ratio. Only putative single units were used for further analyses.

### Maximum Noise Entropy (MNE) Models

We fit MNE stimulus-response functions [43, 44, 37] to the song evoked responses of each neuron assuming each of the acoustic feature representations as the “stimulus”. For signal representations, we pairwise averaged the original spectrogram magnitudes across both frequency and time axes to form 16 × 16 (256) dimensional spectrogram segments, and term the resulting MNEs as signal-CRFs, where CRF refers to the composite receptive field [37] expressing both the linear and nonlinear components of the neuron’s spiking response as a function of the stimulus representation. For the predictive representation, we used the 256-dimension latent space activation matrix obtained by feeding song spectrograms for the physiological test stimuli into the song-trained TCM. For the prediction-error representation we used the pixel-wise squared difference between the TCM predicted future spectrogram segment associated with each segment of the physiological test stimuli and the true future spectrogram segment, applying the same pairwise averaging across both frequency and time to form a 16×16 (256) dimensional spectrogram segment. Stimulus samples were z-scored before inputting into their corresponding MNE models. To maintain temporal alignment between spiking responses and the various stimulus representations, we binned each unit’s spike train in time to correspond with the outputs of the TCM as described above to yield 10.5 msec bins each containing a count of the number of spikes from the neuron during that time interval. We split the spiking responses into 80-20% subsets for MNE training and testing, respectively.

### Fitting MNE models

The MNE model fits a logistic function to capture the probability of spiking in a neuron given a linear combination of first- and second-order features of the stimulus (equation 1). Parameters a, h and J are optimized to constrain stimulus-response relationships corresponding to the mean firing rate, spike-triggered average and spike-triggered covariance respectively. The model optimization process uses a four-fold jackknife approach, where acoustic inputs and response data are split into four batches. In each jackknife iteration, three of the batches are used as training data, and the remaining batch was reserved for testing. Using a nonlinear conjugate gradient algorithm, the log loss between the response and the weighted input was minimized. To prevent overfitting, early stopping with a ten-epoch criterion on the testing data log loss is enforced as a regularization measure. The optimized weights from each jackknife are then averaged, yielding a single set of mean weights used in all subsequent analyses. A detailed description of the method is given in [43]. The optimized MNE model takes the form:

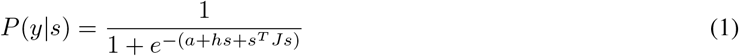

where *P* (*y*|*s*) is a time series of predicted spiking probability conditioned on the stimulus, *s*.

### Modeling MNE responses to novel stimuli

We used trained MNE models to generate predicted neural responses to novel (not used for training) songs. Response predictions were generated using either the complete set of learned parameters (a, h, J) and s, or only the trained linear features (parameters a and h) and s, using equation 1. To assess prediction accuracy, we calculated the Pearson’s correlation coefficient (r) between the predicted spiking probability and the empirical response expressed as the averaged trial-average spike counts for each unit (Figure 2E). These correlation values were used to compare prediction quality across MNE models across different acoustic feature representations. Shuffling spiking responses. To estimate the null distribution of MNE-predicted responses due to chance, we followed the same MNE fitting procedure as above but using permuted versions of spiking responses from each neuron. Specifically, we shuffled the timing of all spikes during a given stimulus presentation, such that the stimulus-locked empirical response is destroyed, while the total number of spikes per stimulus is preserved. To assess the quality of predicitons, we calculated the Pearson’s correlation coefficient (*r*) between the predicted spiking probability from null distribution trained CRFs and the empirical response.

### Partitioning of MNE-modeled response variance

To partition the variance in the response predicted by the CRFs tied to each of the three separate feature spaces, we used a regression analysis adapted from multiple linear regression and custom written in R, in which the empirical response of each unit to song is predicted as various linear combinations of the signal, prediction and prediction-error MNE modeled responses. We fit models containing each feature space (signal, prediction, prediction-error), pairs of features spaces (signal plus prediction, signal plus error, and prediction plus error), and all three feature spaces (signal plus prediction plus error) using a standard linear model and computed an adjusted R-squared score (equation 2) as a measure of explained response variance for each fit. Adjusted R-squared scores range from 0 to 1.

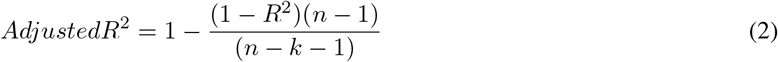

We calculated partial Pearson’s correlation coefficients, treating the empirical neural response as a composition of modeled response probabilities, using the *ppcor* package in *R* [81], and report unique and shared response variance attributable to the signal, prediction and prediction-error representations.

### Functional specialization in single neurons

#### Unit bias metric

We quantified bias in individual units by calculating the unique response variance attributable to prediction-error (e) and prediction (p). A bias metric for each unit was defined as the difference between the partial Pearson’s correlation coefficients for error and prediction (i.e., e minus p). We calculated Spearman’s *r* for each unit’s prediction and error pair by ranking the data and computing the correlation of these ranks. The coefficient ranges from -1 to 1, with values near 1 indicating a strong positive monotonic relationship, near -1 a strong negative relationship, and near 0 no monotonic relationship.

#### Functional specialization counts

To identify functionally specialized units within each region, we first shuffle-corrected the bias metric by generating distributions of bias residuals for each region. These distributions were obtained using the bias metric calculated from the partial Pearson’s correlation coefficients of the full model MNE-predicted responses and from the null distribution (generated from shuffled spike trains as described above, Figure S7). Units whose empirical bias metric fell within the 95% confidence interval (CI; or margin of error) with respect to the null distribution were not considered. The margin of error was determined using the standard deviation of the corresponding bias residual distribution. We then counted specialized units as those whose bias metric lie in the tails (relative to the 95% CI) of the shuffle-corrected bias residual distribution. Units located in the right-sided tail of the distribution were considered specialized for prediction-error (*e*), while those in the left-sided tail were considered specialized for prediction (*p*) (Table S5).

#### Normality of bias residuals distribution

To assess the normality and characteristics of the bias residual distributions, we employed quantile-quantile (Q-Q) plots to compare bias residuals against the null bias residual distribution for each region (Figure 5). In Q-Q plots, the quantiles of the empirical bias distribution (plotted on the y-axis) are compared to the quantiles of the reference or null distribution (plotted on the x-axis). Interpretation of these plots hinges on the relative positions of the data points. If the data points lie closely along the 45-degree reference line, the tested data distribution matches the reference distribution, indicating similar quantiles and no significant deviation in shape, skewness, or spread. Points below the line suggest a left-skewed distribution, with a longer left tail, while points above the line indicate a right-skewed distribution. A noticeable curvature at the tails indicates skewness: dipping below at lower quantiles and above at higher quantiles suggests left-skewness, and the opposite pattern indicates right-skewness. We performed statistical tests (e.g., Shapiro-Wilk, skewness, and kurtosis, Table S6) to complement Q-Q plots for more formal assessments of normality or distributional characteristics [82]. Shapiro-Wilk test assesses whether a dataset follows a normal distribution, with a significant result (p < 0.05) indicating deviation from normality. Skewness measures the asymmetry of a distribution: a value of 0 indicates perfect symmetry, positive skewness (> 0) suggests a right-skewed distribution (longer right tail), and negative skewness (< 0) indicates left-skewness (longer left tail).

Fisher’s (or excess) kurtosis quantifies the “tailedness” of the distribution, where high kurtosis (> 0) indicates heavy tails (more extreme values), and low kurtosis (< 0) suggests lighter tails or a flatter distribution. Together, these metrics provide insights into deviations from normality in data.

### Clustering of unique variances

We used HDBSCAN [49] (*scikit-learn*) to cluster the partial Pearson’s correlation coefficients of signal, prediction and prediction-error of individual units within each simultaneously recorded neural population. Each clustering used default parameterization of HDBSCAN, setting the minimum samples at 1. We varied the minimum cluster size parameter to achieve the best possible clusterability as quantified by silhouette score (Equation 3 and 4). The reported silhouette scores are the mean silhouette coefficient across all the samples in a dataset. The silhouette coefficient measures how distant each point is to points in its own category, relative to its distance from the nearest point in another category. It is therefore taken as a measure of how tightly clustered elements are that belong to the same category. Silhouette scores range from -1 to 1, with 1 being more clustered. The silhouette score S, is computed as the mean of the silhouette coefficients for each data point. For each data point (i), the silhouette coefficient si is the mean distance between the data point and all other data points in the same cluster (ai), minus the distance to that point’s nearest neighbor belonging to a different cluster (bi), divided by the maximum of ai and bi, written as:

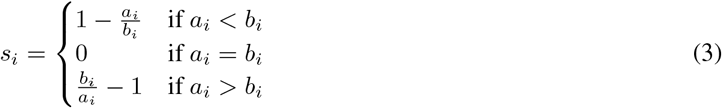

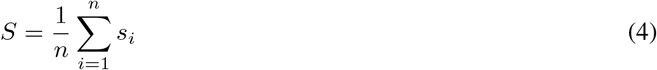

### Statistical analyses

All statistical analyses were performed with Python or R. Non-parametric statistical tests were applied in cases where data samples violated the assumptions required for parametric tests, such as normality or homoscedasticity. Unless otherwise noted, we express all means with the corresponding standard errors. We utilized Linear Mixed Effects (LME) models to analyze the data, which account for both fixed and random effects (e.g., recording blocks or subject), allowing us to model individual variability and account for correlations within grouped data. Effect sizes were estimated with Cliff’s delta for samples that did not conform to a normal distribution. We used the standard p-value < 0.05 for assigning statistical significance denoted by asterisk symbol. In cases where the sample size was large (n > 25), the standard p-value was used in conjunction with a Cliff’s delta > 0.33, which are traditionally assigned to a medium-sized effects or greater. We used t-tests and paired t-tests for post-hoc comparisons.

## Supplemental Information

**Table S1:** Temporal Convolutional Model (TCM) architecture outline. We employed a convolutional architecture featuring a 256-dimensional latent space. The encoder consists of five convolutional layers followed by a fully connected layer, while the decoder mirrors this structure with a fully connected layer and five deconvolutional layers. Leaky ReLU was applied as the activation function for artificial neurons. The network was trained using batches of 128 segments. The training process utilized the ADAM optimizer in TensorFlow.

| Encoder |  | Decoder |  |
| --- | --- | --- | --- |
| layer | architecture | layer | architecture |
| input | (128, 32, 32, 1) | input | (128, 256) |
| convolutional | (128, 30, 30, 8) | fully connected | (128, 1024) |
| convolutional | (128, 28, 28, 16) | Reshape | (128, 2, 2, 256) |
| convolutional | (128, 26, 26, 32) | deconvolutional | (128, 4, 4, 64) |
| convolutional | (128, 24, 24, 64) | deconvolutional | (128, 8, 8, 64) |
| convolutional | (128, 22, 22, 64) | deconvolutional | (128, 16, 16, 32) |
| fully connected | (128, 30976) | deconvolutional | (128, 32, 32, 16) |
| fully connected | (128, 256) | deconvolutional | (128, 32, 32, 1) |

**Table S2:** Neural datasets. We recorded from 8 (NCM, n = 3; CMM, n = 4; Field L, n = 3) subjects over 14 blocks (NCM, n = 3; CMM, n = 5; Field L, n = 6) of recordings. Acute implanted subjects performed 20 trials of each of the 60 songs in the stimulus set during recording. The silence interval between trials was randomly sampled between 2 and 5 seconds. Recording blocks were obtained from independent populations of cells in both hemispheres and at different depths within the region of interest.

| Subject | Recording block | Probe location | Brain region | Hemisphere | Number of units |
| --- | --- | --- | --- | --- | --- |
| **B1240** | 1 | AP:-125; ML:1125; DV:2100 | NCM | Right | 214 |
| **B1596** | 1 | AP:250; ML:750; DV:2900 | NCM | Right | 160 |
| **B1692** | 1 | AP:0; ML:1500; DV:1445 | NCM | Right | 194 |
| **B1425** | 1 | AP:2500; ML:500; DV:2320 | CMM | Right | 151 |
| **B1260** | 1 | AP:2450; ML:300; DV:2700 | CMM | Right | 172 |
| **B1152** | 1 | AP:2500; ML:500; DV:2530 | CMM | Right | 82 |
| **B1152** | 2 | AP:1800; ML:1800; DV:1800 | Field L | Right | 93 |
| **B1584** | 1 | AP:2000; ML:1900; DV:2350 | Field L | Right | 154 |
| **B1584** | 2 | AP:1800; ML:1800; DV:1710 | Field L | Left | 96 |
| **B1584** | 3 | AP:1800; ML:1800; DV:3110 | Field L | Left | 159 |
| **B1457** | 1 | AP:1800; ML:1800; DV:1850 | Field L | Left | 80 |
| **B1457** | 2 | AP:1800; ML:1800; DV:1200 | Field L | Left | 40 |
| **B1457** | 3 | AP:2450; ML:500; DV:2750 | CMM | Right | 69 |
| **B1457** | 4 | AP:2450; ML:500; DV:2200 | CMM | Right | 85 |

**Table S3:** Comparisons of song- and MNIST-trained TCM networks. Mean (± SEM) Pearson’s correlation coefficients for each region showing relationship between the empirical response to song and the response predicted by composite receptive fields (CRFs) fit to predictive and prediction-error features obtained from TCM network trained on song (left) or MNIST digits (right), with associated p-values and t-statistics for the paired comparison (neuron to neuron).

| Region | Song prediction-CRF | Song error-CRF | MNIST prediction-CRF | MNIST error-CRF |
| --- | --- | --- | --- | --- |
| NCM | 0.76 ± 0.004 | 0.48 ± 0.004 | 0.55 ± 0.005<br>t = -97.0, p = 0 | 0.53 ± 0.004<br>t = 23.9, p = 4.4e-88 |
| CMM | 0.79 ± 0.005 | 0.64 ± 0.008 | 0.65 ± 0.007<br>t = -29.0, p = 2.4e-113 | 0.66 ± 0.007<br>t = 9.4, p = 1.4e-19 |
| Field L | 0.74 ± 0.007 | 0.70 ± 0.008 | 0.63 ± 0.009<br>t = -18.8, p = 1.1e-62 | 0.70 ± 0.007<br>t = 3.0, p = 0.003 |

**Table S4:** Partial Pearson’s correlation coefficients by region. Partial Pearson’s correlation coefficients were calculated treating the empirical neural response as a composition of modeled response probabilities were used to report unique and shared variances of signal, prediction and prediction error. Table shows Partial Pearson’s correlation coefficients for modeled responses computed using either the complete MNE or shuffled MNE model across NCM, CMM and Field L, along with relevant Kolmogorov-Smirnov test statistic and associated p-value comparing the empirical and shuffled distributions.

| Region | Variance Term | Full MNE | Shuffled MNE | KS statistic; p value |
| --- | --- | --- | --- | --- |
| NCM | signal | 0.33 ± 0.005 | 0.00 ± 0.001 | statistic = 0.98, p = 8.23e-313 |
| NCM | prediction | 0.39 ± 0.004 | 0.00 ± 0.001 | statistic = 0.99, p = 0.0 |
| NCM | error | 0.11 ± 0.002 | 0.00 ± 0.000 | statistic = 0.92, p = 4.00e-261 |
| NCM | prediction + error | 0.1 ± 0.002 | 0.02 ± 0.001 | statistic = 0.77, p = 5.99e-165 |
| NCM | signal + error | 0.16 ± 0.003 | 0.02 ± 0.001 | statistic = 0.94, p = 1.42e-275 |
| NCM | prediction + signal | 0.51 ± 0.004 | 0.14 ± 0.003 | statistic = 0.91, p = 2.60e-250 |
| NCM | signal + prediction + error | 0.04 ± 0.001 | 0.00 ± 0.000 | statistic = 0.96, p = 5.26e-293 |
| NCM | pred. + error (minus shared) | 0.06 ± 0.003 | 0.02 ± 0.000 |  |
| NCM | signal + error (minus shared) | 0.12 ± 0.002 | 0.02 ± 0.000 |  |
| NCM | pred. + signal (minus shared) | 0.47 ± 0.004 | 0.14 ± 0.003 |  |
| CMM | signal | 0.17 ± 0.006 | 0.00 ± 0.001 | statistic = 0.74, p = 2.79e-148 |
| CMM | prediction | 0.29 ± 0.008 | 0.00 ± 0.001 | statistic = 0.84, p = 4.42e-202 |
| CMM | error | 0.06 ± 0.003 | 0.00 ± 0.001 | statistic = 0.46, p = 5.03e-53 |
| CMM | prediction + error | 0.22 ± 0.005 | 0.10 ± 0.002 | statistic = 0.60, p = 4.89e-93 |
| CMM | signal + error | 0.27 ± 0.006 | 0.10 ± 0.003 | statistic = 0.58, p = 4.09e-86 |
| CMM | prediction + signal | 0.63 ± 0.008 | 0.24 ± 0.005 | statistic = 0.78, p = 1.07e-164 |
| CMM | signal + prediction + error | 0.12 ± 0.005 | 0.02 ± 0.001 | statistic = 0.64, p = 1.16e-105 |
| CMM | pred. + error (minus shared) | 0.10 ± 0.004 | 0.08 ± 0.002 |  |
| CMM | signal + error (minus shared) | 0.15 ± 0.003 | 0.08 ± 0.002 |  |
| CMM | pred. + signal (minus shared) | 0.51 ± 0.011 | 0.22 ± 0.004 |  |
| Field L | signal | 0.09 ± 0.006 | 0.00 ± 0.001 | statistic = 0.44, p = 3.81e-55 |
| Field L | prediction | 0.23 ± 0.007 | 0.01 ± 0.001 | statistic = 0.76, p = 3.10e-176 |
| Field L | error | 0.05 ± 0.002 | 0.00 ± 0.001 | statistic = 0.55, p = 4.77e-86 |
| Field L | prediction + error | 0.29 ± 0.005 | 0.13 ± 0.002 | statistic = 0.65, p = 1.28e-123 |
| Field L | signal + error | 0.34 ± 0.004 | 0.11 ± 0.002 | statistic = 0.80, p = 6.94e-200 |
| Field L | prediction + signal | 0.66 ± 0.007 | 0.27 ± 0.005 | statistic = 0.81, p = 1.17e-211 |
| Field L | signal + prediction + error | 0.18 ± 0.004 | 0.03 ± 0.001 | statistic = 0.87, p = 2.01e-247 |
| Field L | pred. + error (minus shared) | 0.11 ± 0.004 | 0.10 ± 0.001 |  |
| Field L | signal + error (minus shared) | 0.16 ± 0.004 | 0.08 ± 0.002 |  |
| Field L | pred. + signal (minus shared) | 0.48 ± 0.010 | 0.24 ± 0.004 |  |

**Table S5:** Functional specialization counts. The number of neurons expressing extreme values relative to the overall distribution of bias metric scores in each region, relative to a threshold set at either the 90% or 95% confidence interval. The bias metric is the difference between the partial correlation coefficients associated with the CRFs fit to the error and predictive representations for each neuron. Parenthetical value gives the same count for distribution of bias metrics computed from the shuffled CRFs.

| Region | Error-biased units |  | Prediction-biased units |  |
| --- | --- | --- | --- | --- |
|  | Rel. 90% CI | Rel. to 95% CI | Rel. to 90% CI | Rel. to 95% CI |
| NCM | 25 (32) | 17 (18) | 12 (30) | 8 (16) |
| CMM | 25 (35) | 11 (17) | 19 (18) | 2 (11) |
| Field L | 7 (20) | 2 (13) | 53 (32) | 29 (17) |

**Table S6:** Statistics for bias metric distributions across all regions. Shapiro-Wilk test statistics assessing whether the distribution of prediction vs. error bias metric values (Figure S8B) is normally distributed. P-values for each region indicate significant deviation from normality. NCM shows a strong positive skew and high kurtosis, indicating more neurons with a bias for predictive over error encoding than expected by chance. Field L is skewed in the opposite direction, indicating a bias for error over predictive coding neurons, though with fewer extremes (low kurtosis) than in NCM. The distribution of CMM bias scores is largely symmetric and non-kurtotic.

| Region | Shapiro-Wilk test | Skew | Kurtosis |
| --- | --- | --- | --- |
| NCM | statistic = 0.94, p = 9.54e-15 | 1.151 | 5.717 |
| CMM | statistic = 0.98, p = 1.05e-05 | 0.096 | -0.800 |
| Field L | statistic = 0.96, p = 7.29e-12 | -0.554 | -0.463 |

**Figure S1:**
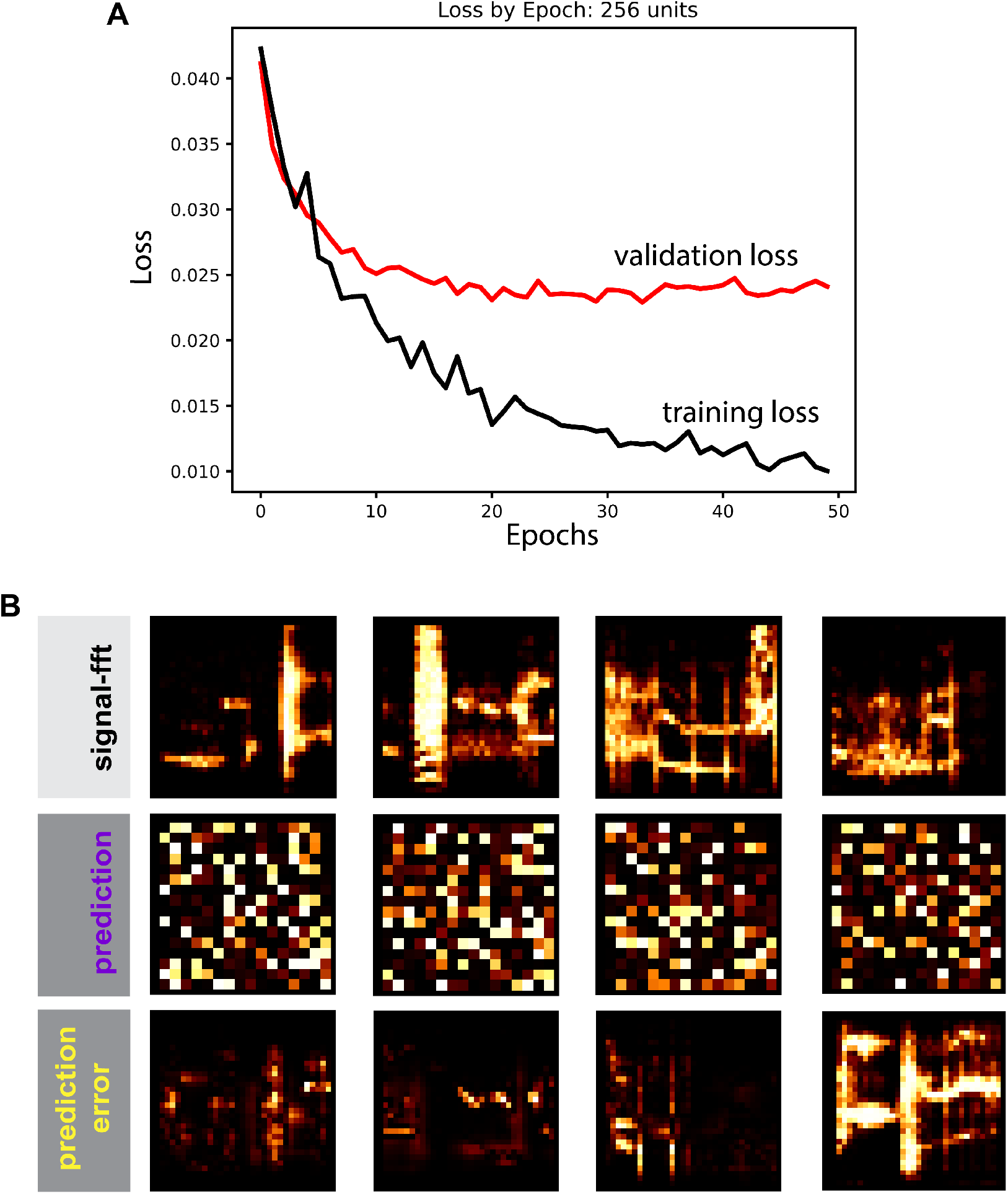
Temporal Convolutional Model (TCM) performance and outputs. A) Temporal Convolutional Model (TCM) performance on held-out validation data comprising 10% of the full dataset not used for training. The plot shows both validation loss (red) and training loss (black) (Methods) over 50 epochs. B) Example acoustic feature representations: signal (top row) comprising general spectrotemporal magnitude coefficients for typical song segments (columns), and the corresponding TCM prediction (middle row) comprising the latent feature weights predictive of upcoming song segments in the TCM network, and prediction-error segments comprising error features obtained from the difference between true spectrotemporal segments and TCM-predicted output. Signal and prediction-error representations are in the spectrographic time-frequency coordinates, prediction representations are in arbitrary axes.

**Figure S2:**
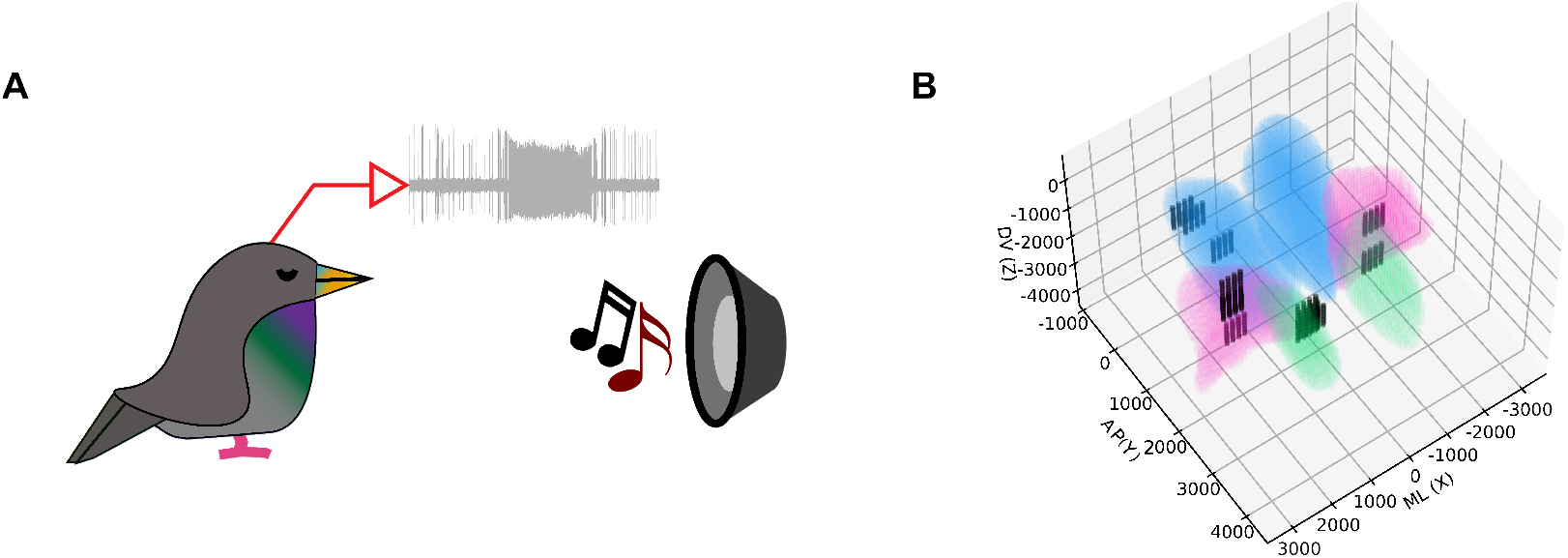
Electrophysiology. A) Extracellular voltage waveforms were collected from multisite silicon probes implanted in NCM, CMM and Field L (Methods) from anesthetized animals while listening to a set of natural birdsongs comprising 1-minute-long songs and seconds-long motif sequences from conspecific singers. B) Visualization of recording sites shown over top of the starling brain atlas showing NCM (blue), CMM (green) and Field L (pink).

**Figure S3:**
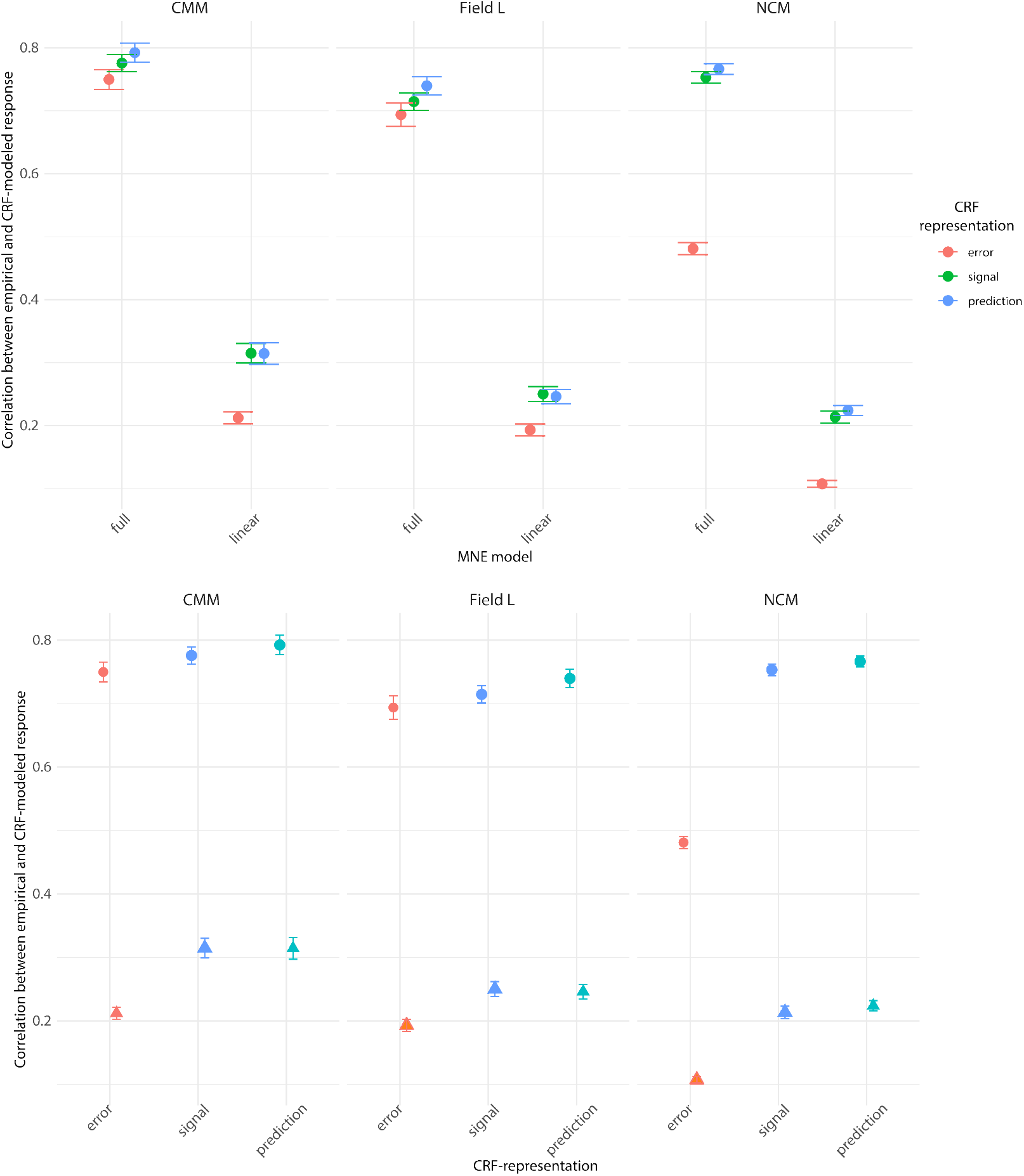
Linear and non-linear MNE model comparisons. Three-way interaction plots showing the complex relationship between stimulus representation (signal, prediction and prediction-error), brain region (CMM, Field L, and NCM) and MNE models (full or linear model). Main effects of representation, region, and model and interactions are described in the text. The three-way interaction is significant (F(4) = 60.018, p < 0.0001). Each plot shows the correlation between the response predicted by a specific CRF-model and the empirical response to novel songs. The top panel highlights the differences between full and linear MNE models fit to the predictive (blue), signal (green), and prediction-error (red) representations. The bottom panel shows the same data broken out according the MNE model, either linear only (triangles) or the full model (circles) which includes both linear and non-linear parameters. Note that the full models outperform the linear models, but the extent of this performance difference varies depending on the brain region and the stimulus representation. In CMM and Field L the full nonlinear models uniformly increase performance over the linear models for all three stimulus representations. In NCM, however, the nonlinear models provide a much smaller benefit to error encoding.

**Figure S4:**
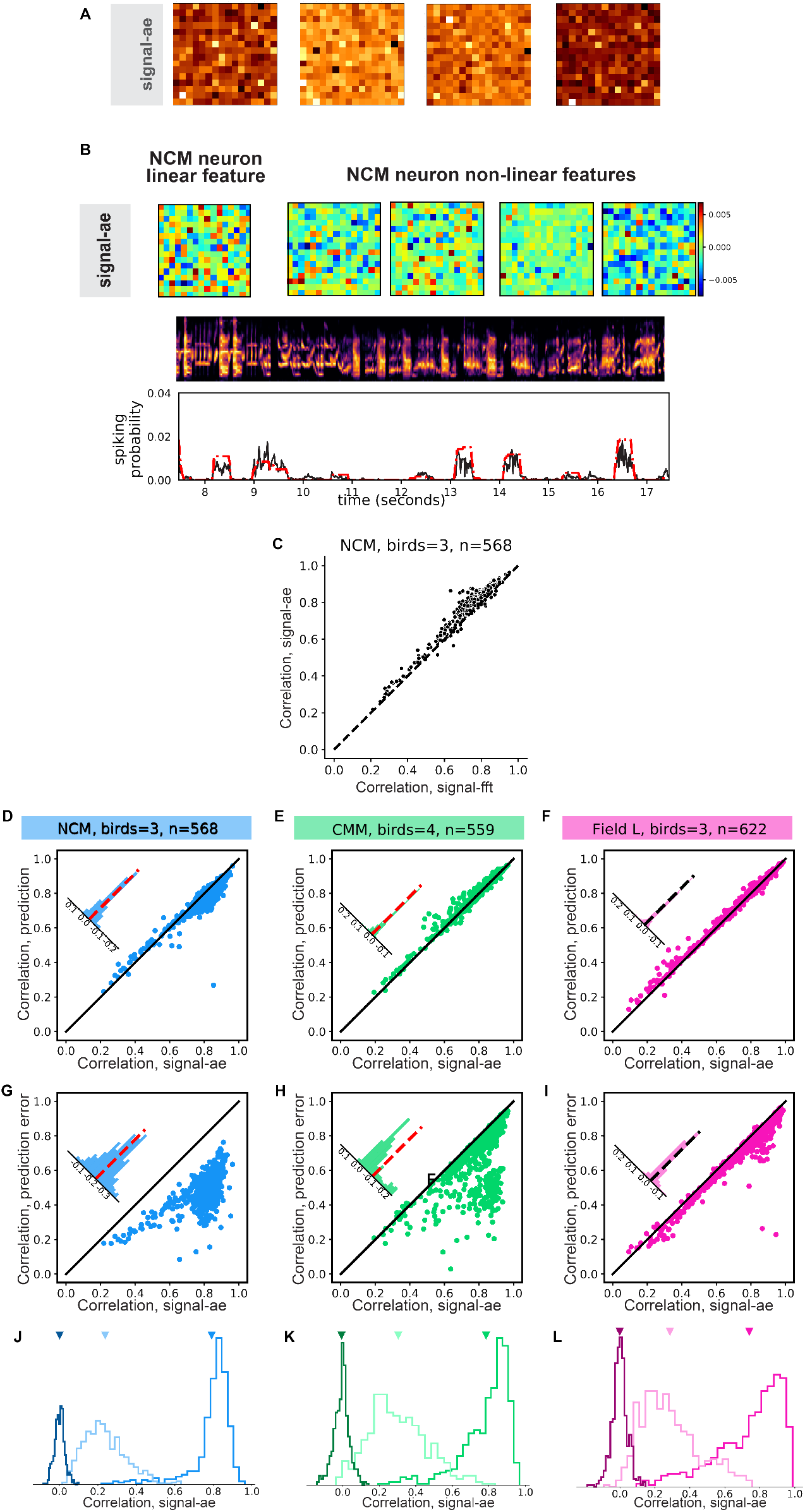
Alternative signal representations. A) Example signal-ae representations for song segments (as in Figures 2D and S1) showing the 256-dimensional latent space activation of an autoencoder network similar to the TCM, but trained to reconstruct the input song segment rather than the future song segment. B) Composite receptive fields (CRF; top) of one NCM neuron fit to signal-ae stimulus representation showing linear component (left, obtained from parameter h of the MNE model) and four most significant non-linear components (right, obtained from MNE parameter J). Note that autoencoder latent weight matrices and their corresponding CRFs do not (Figure S4 continued) have a 2-dimensional spectro-temporal structure. Empirical response and ae-CRF-predicted response predictions (bottom) to a novel stimulus (middle, spectrogram) not used to compute the CRF (r(model,empirical)=0.80). C) Scatterplot of response correlations between CRF modeled and empirical2n7eural responses using CRFs trained on either spectrographic (signal-CRF) or autoencoder latent space (ae-CRF) representations of the acoustic stimulus. Each point corresponds to a single unit in NCM (n=568). Scatterplots showing the comparative performance (correlation between empirical and predicted response) for CRF models fit to the signal-ae and the TCM prediction representations of song in (D) NCM, (E) CMM and (F) Field L, and for CRF models fit to the signal-ae and the TCM prediction-error representations of song in the same regions (G-I). Dashed red lines in inset residual histograms indicate the position of mean in each distribution, the diagonal line indicates unity. (J-K) Distributions of correlations between CRF-predicted and empirical responses in (J) NCM (n=568 neurons), (K) CMM (n=559), and (L) Field L (n=622). Dark colors show correlations for shuffled control models (Methods), medium colors for full linear/non-linear model, and light colors to linear only model. Arrows indicate the position of means in each distribution.

**Figure S5:**
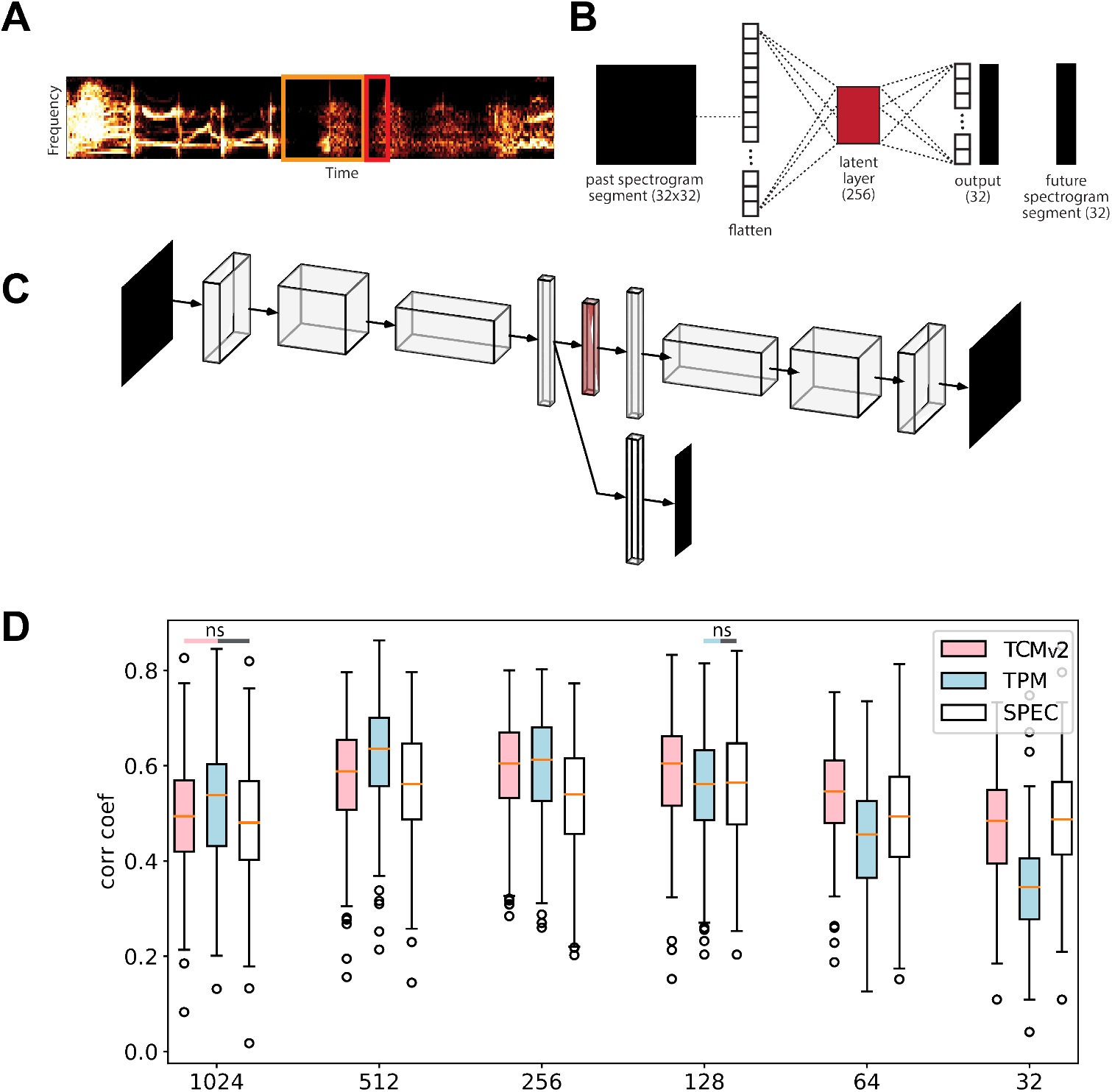
Alternative generative model architectures and TCM optimization. An alternate spectrogram segmentation scheme (A; compare to main text and Figure 1) and predictive network model architecture (B), referred to as the Temporal Prediction Model (TPMv1), in which past spectrogram segments (32 frequency bins x 32 time bins) predict future segments of only one time bin in length (32 frequency bins x 1 time bin) using a single linear feedforward layer and nonlinear sigmoidal activation. (C) An alternate Temporal Convolutional Model architecture (TCMv2) in with parallel decoders for prediction and reconstruction. Past segments of 32×32 dimensions are used to reconstruct the full (32 x32) input (top) and to predict the next immediate time bin (1×32, bottom). (D) Box plots showing Pearson’s correlation coefficients, r, between the CRF-predicted and empirical responses, computed from network latent space or spectrographic representations of differing sizes from 32 – 1024 dimensions. The spectrogram representations (white) were pairwise averaged to match the dimensionality of the latent space in the TCMv2 (pink) and TPM (blue) networks.

**Figure S6:**
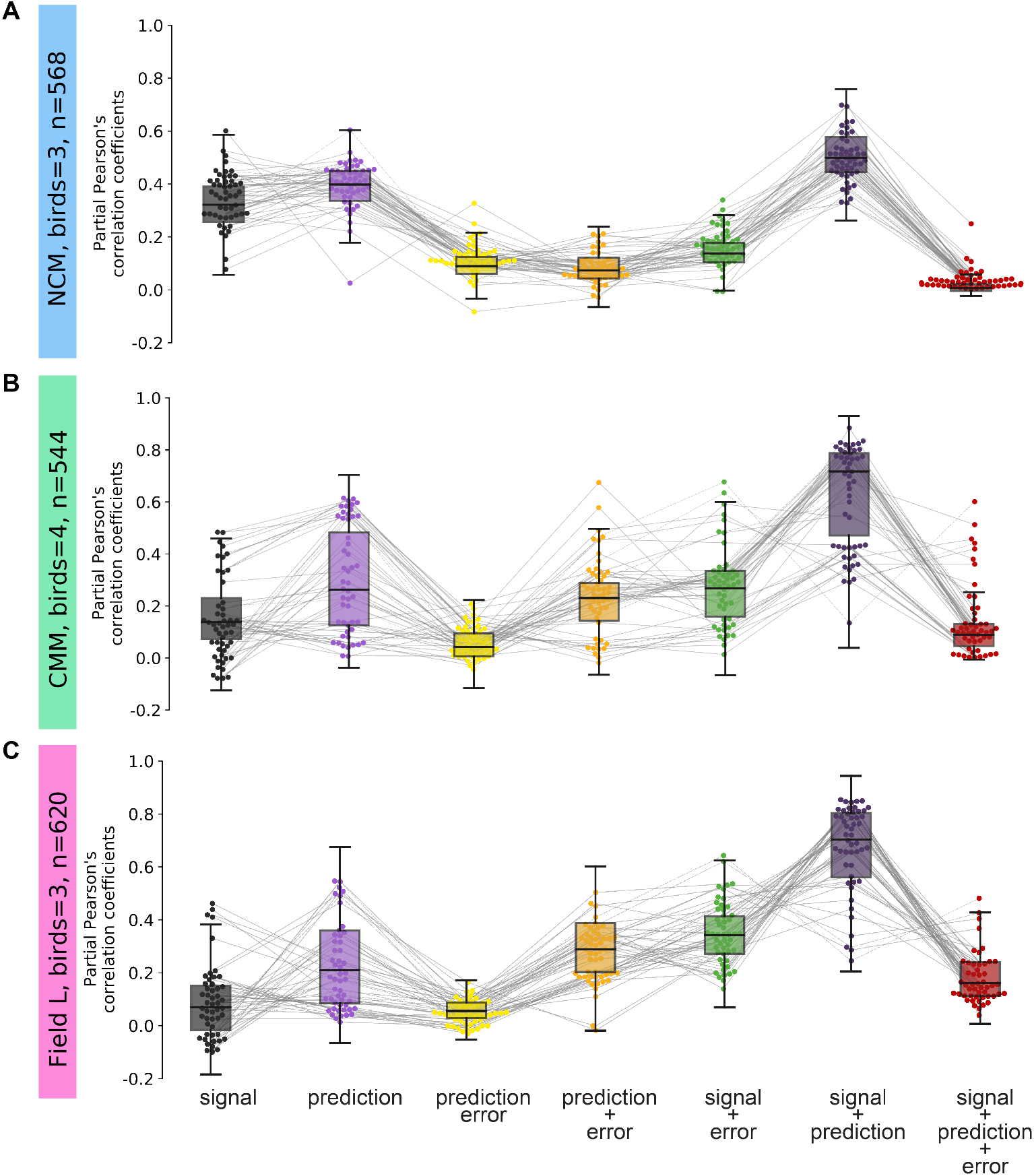
Unique and shared variance partitions for signal, prediction and prediction-error representations. Box plots showing distribution of the partial Pearson’s correlation coefficients for the unique contributions of signal, prediction, prediction-error representations in (A) NCM (n=568 neurons), (B) CMM (n=544), and (C) Field L (n=620). Boxplots represent the overall distribution of all neurons in each region, overlaid points show a random sample of 50 neurons in each region. Two-way analysis of variance (ANOVA), main effect for stimulus representation, *F* = 1522.6, *df* = 2, *p* < 2.2 − *e*16, main effect for brain region, *F* = 743.9, *df* = 2, *p* < 2.2 − *e*16, stimulus representation × brain region interaction term, *F* = 88.1, *df* = 4, *p* < 2.2 − *e*16.

**Figure S7:**
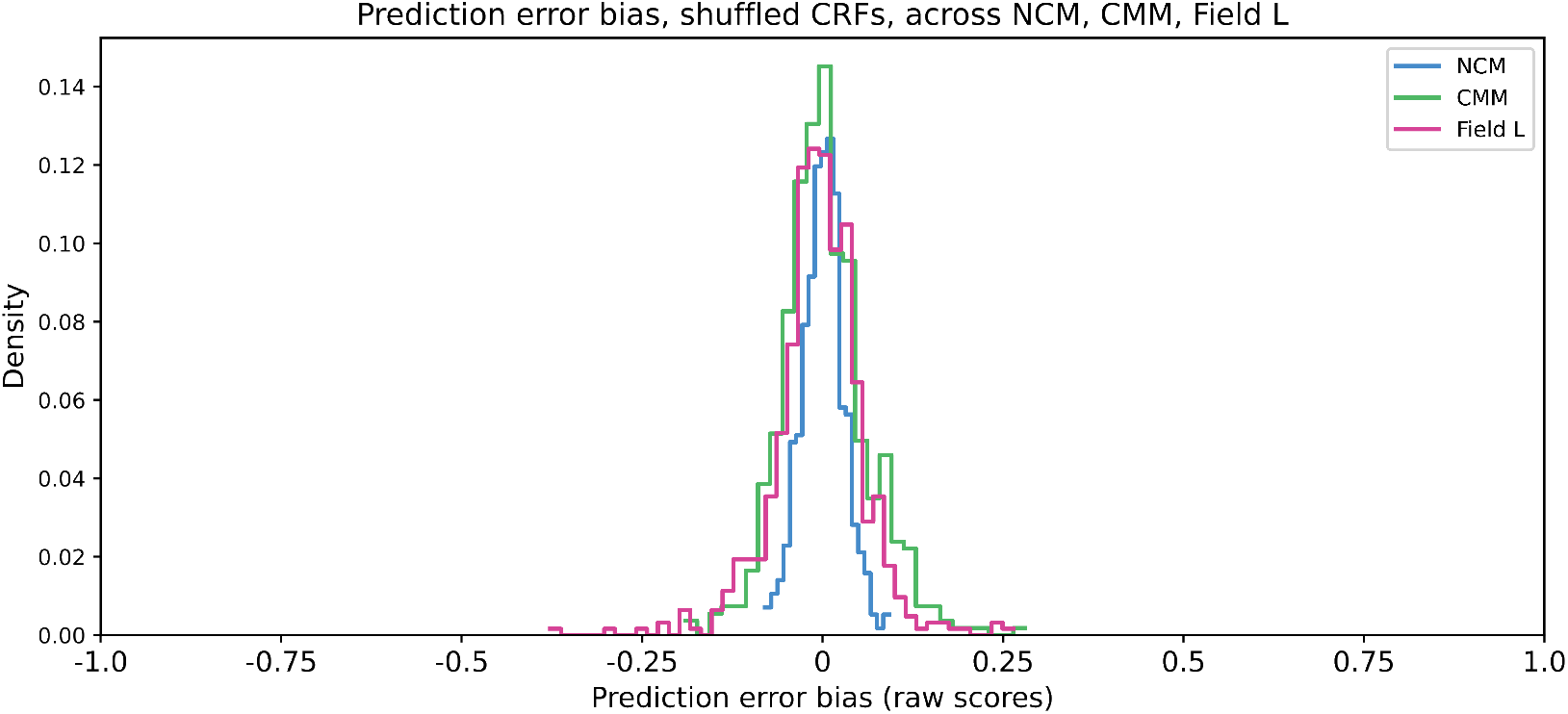
Shuffle control distributions of prediction error bias in single neurons. Histograms showing distribution of bias residuals, determined by the difference between unique error and unique prediction (error - prediction), in NCM, CMM and Field L computed from the output of CRFs fit to the shuffled spike trains for neurons in each region (mean ± SEM NCM = 0.001±0.001; CMM = −0.004±0.002; Field L = −0.006±0.002). Differences between the respective empirical (Figure 5) and shuffled distribution are significant (NCM, statistic = 0.977, *p* = 7.12*e* − 311; CMM, statistic = 0.742, *p* = 1.45*e* − 143; Field L, statistic = 0.600, *p* = 1.83*e* − 101; two-sample Kolmogorov-Smirnov test).

**Figure S8:**
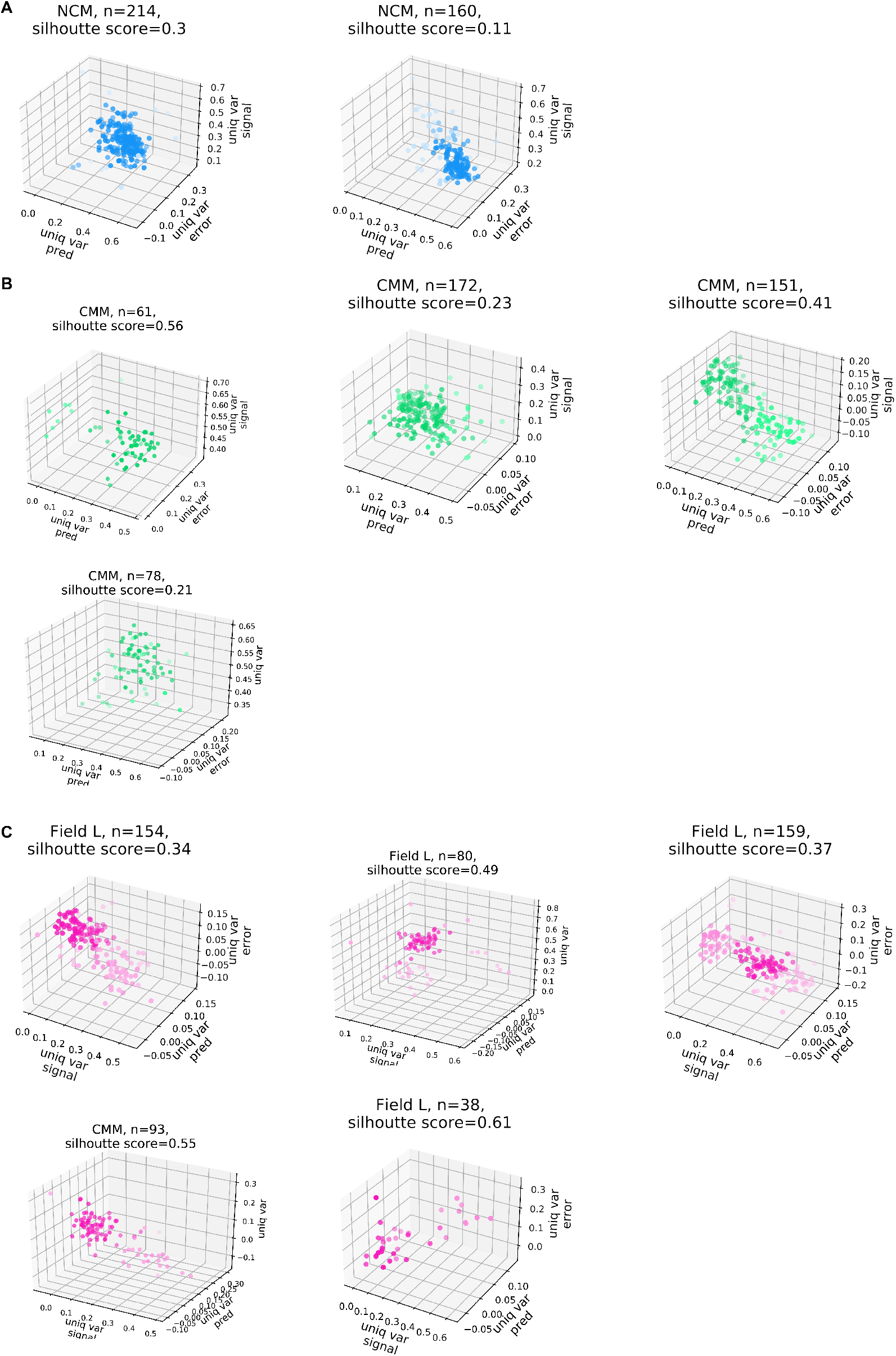
Clustering of auditory neurons based on signal, prediction and prediction-error responses. Three-dimensional scatterplots showing partial Pearson’s correlation coefficients attributable to signal, prediction, and error representations in for populations of simultaneously recorded neurons in (A) NCM (blue) (B) CMM (green) and (C) Field L (pink). Clusterability measure is determined by silhouette score from HDBSCAN (Methods).

**Figure S9:**
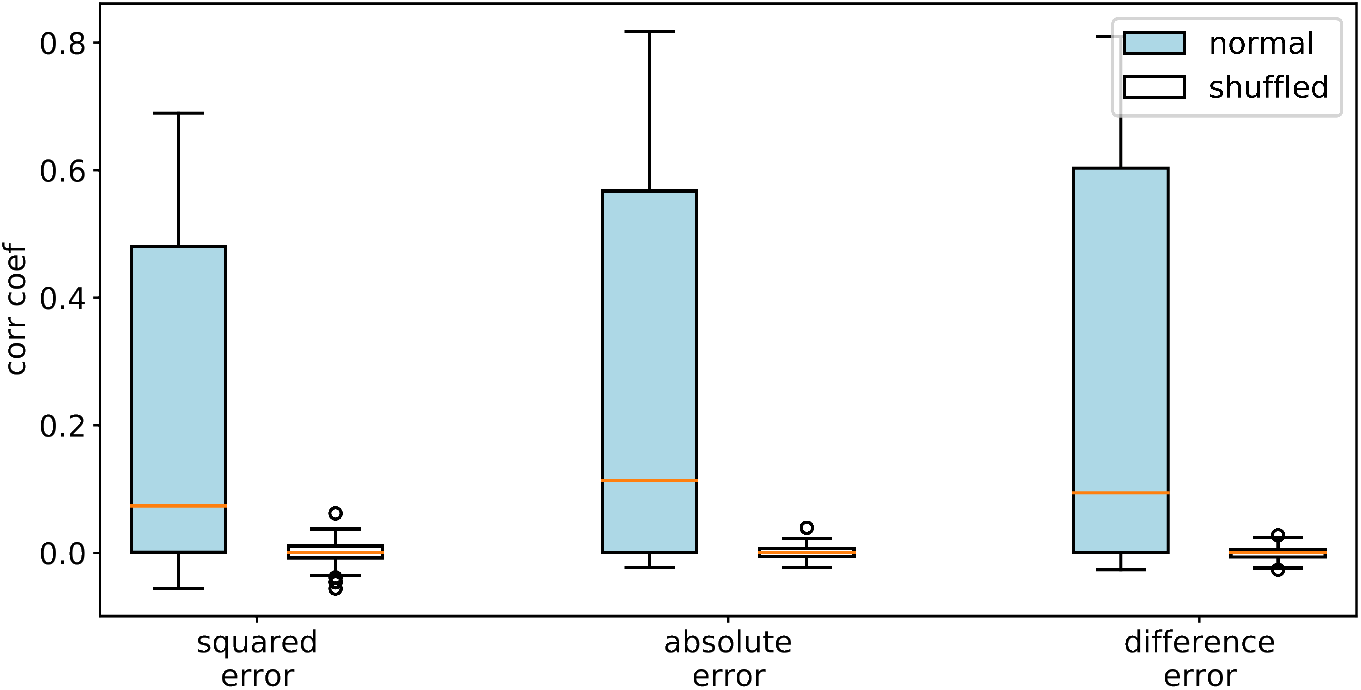
Alternate prediction-error estimates yield similar results. Correlation coefficients between empirical responses and the CRF predicted response with the CRF fit to three different estimates of the prediction-error computed using either the squared difference between the TCM-predicted song and the true song, (*a* − *b*)^2^, the absolute value of the difference, |*a* − *b*|, or the difference (*a* − *b*). For comparison in each case, the Pearson’s correlation coefficients (*r*) for full MNE models fit to both the empirical spiking response and a shuffled control, obtained by shuffling stimulus segments disrupting stimulus and response locking. All three measures of prediction error produced significant results (higher response encoding) compared to chance (squared error: *t* = 78.60, *p* = 2.63*e* − 159; absolute error: *t* = 94.30, *p* = 1.13*e* − 175; difference error: *t* = 109.15, *p* = 6.18*e* − 18, paired t-test).

